# The defining genomic and predicted metabolic features of the Acetobacterium genus

**DOI:** 10.1101/2020.01.23.904417

**Authors:** Daniel E. Ross, Christopher W. Marshall, Djuna Gulliver, Harold D. May, R. Sean Norman

## Abstract

Acetogens are anaerobic bacteria capable of fixing CO_2_ or CO to produce acetyl-CoA and ultimately acetate using the Wood-Ljungdahl pathway (WLP). This autotrophic metabolism plays a major role in the global carbon cycle. *Acetobacterium woodii*, which is a member of the *Eubacteriaceae* family and type strain of the *Acetobacterium* genus, has been critical for understanding the biochemistry and energy conservation in acetogens. Other members of the *Acetobacterium* genus have been isolated from a variety of environments or have had genomes recovered from metagenome data, but no systematic investigation has been done into the unique and varying metabolisms of the genus. Using the 4 sequenced isolates and 5 metagenome-assembled genomes available, we sequenced the genomes of an additional 4 isolates (*A. fimetarium, A. malicum, A. paludosum,* and *A. tundrae*) and conducted a comparative genome analysis of 13 different *Acetobacterium* genomes to obtain better phylogenomic resolution and understand the metabolic diversity of the *Acetobacterium* genus. Our findings suggest that outside of the reductive acetyl-CoA (Wood-Ljungdahl) pathway, the *Acetobacterium* genus is more phylogenetically and metabolically diverse than expected, with metabolism of fructose, lactate, and H_2_:CO_2_ constant across the genus, and ethanol, methanol, caffeate, and 2,3-butanediol varying across the genus. While the gene arrangement and predicted proteins of the methyl (Cluster II) and carbonyl (Cluster III) branches of the Wood Ljungdahl pathway are highly conserved across all sequenced *Acetobacterium* genomes, Cluster 1, encoding the formate dehydrogenase, is not. Furthermore, the accessory WLP components, including the Rnf cluster and electron bifurcating hydrogenase, were also well conserved, though all but four strains encode for two Rnf clusters. Additionally, comparative genomics revealed clade-specific potential functional capabilities, such as amino acid transport and metabolism in the psychrophilic group, and biofilm formation in the *A. wieringae* clade, which may afford these groups an advantage in low-temperature growth or attachment to solid surfaces, respectively. Overall, the data presented herein provides a framework for examining the ecology and evolution of the *Acetobacterium* genus and highlights the potential of these species as a source of fuels and chemicals from CO_2_-feedstocks.

## Introduction

Acetogens are ubiquitous in nature, phylogenetically diverse, and produce acetyl-CoA from fixation of two molecules of CO_2_, producing acetate as the sole product from the reductive acetyl-CoA/Wood-Ljungdahl pathway (WLP) [1]). Of the over 22 genera that contain acetogens, *Clostridium* and *Acetobacterium* contain the most acetogenic species [2]. *Acetobacterium* spp. are of particular interest because the WLP is the genus’ defining feature and *A. woodii* has been extensively studied. The genus *Acetobacterium* contains gram-positive, non-spore-forming, homoacetogenic bacteria, and was first described as a genus by Balch and coworkers, with the type strain *Acetobacterium woodii* WB1 (ATCC 29683) [3]. Members of the genus *Acetobacterium* have been found in diverse environments, including sulfate-reducing permeable reactive zones [4], anoxic bottom waters of a volcanic sub-glacial lake [5, 6], seagrass rhizosphere [7], high-temperature gas-petroleum reservoirs [8], anaerobic granular sludge from fruit-processing wastewater [9] and biocathode communities [10–14]. Sulfate reducing bacteria (SRB) are often found in these environments as well, with several studies suggesting a syntrophic partnership between SRB and acetogens [15, 16]. In particular, a combination of *Desulfovibrio, Sulfurospirillum,* and *Acetobacterium* were proposed to cooperatively participate in microbial induced corrosion (MIC) of steel [16], have been found in production waters from a biodegraded oil reservoir [17], and can be detected in natural subsurface CO_2_ reservoirs [18]. Moreover, these three microorganisms were the most abundant members of the biocathode community responsible for electrode-driven production of acetate and hydrogen from CO_2_ [19–21].

To date, fourteen sequenced *Acetobacterium* genomes are publicly available— one complete genome of *Acetobacterium woodii* [22], and thirteen draft genomes (including metagenome-assembled genomes) of varying completeness (Table 1). *Acetobacterium woodii* is the type strain of the genus, is the best characterized strain, and has been used as a model organism for understanding the bioenergetics of acetogens ([23], and references therein). Importantly, a genetic system also exists for *A. woodii* [24, 25]. While *A. woodii* utilizes the Wood-Ljungdahl Pathway (WLP) for autotrophic growth, alternative pathways for carbon utilization are operative. Alternative pathways include glucose metabolism, 1,2-propanediol degradation, 2,3-butanediol oxidation, ethanol oxidation, caffeate reduction, ethylene glycol metabolism, and alanine metabolism [26–31].

**Table 1.** Genome characteristics of 14 *Acetobacterium* isolates and MAGs.

|  | Genome size | GC% | Type of genome | Number of contigs | Completeness | Contamination | Source | Reference |
| --- | --- | --- | --- | --- | --- | --- | --- | --- |
| ***A. woodii***<br>(CP002987.1) | 4,044,777 | 39.3 | isolate | 1 | 100 | 1.43 | black sediment | [3, 22] |
| ***A. sp. KB-1***<br>(CP030040.1) | 3,976,829 | 42.5 | MAG | 1006 | 100 | 0.71 | anaerobic bioreactor | Tang et al., 2018, unpublished |
| ***A. bakii***<br>(LGYO00000000.1) | 4,135,228 | 41.2 | isolate | 103 | 99.29 | 2.86 | paper mill waste water pond | [105, 106] |
| ***A. dehalogenans***<br>(AXAC00000000.1) | 4,045,770 | 43.8 | isolate | 81 | 99.29 | 0 | wastewater sludge | [107] |
| ***A. sp. MES1***<br>(MJUY00000000.1) | 3,654,423 | 44.3 | MAG | 65 | 99.29 | 0 | brewery wastewater | [36, 108] |
| ***A. malicum* DSM 4132<sup>T</sup>**<br>(WJBE00000000) | 4,086,418 | 43.7 | isolate | 92 | 99.29 | 1.43 | freshwater sediment | [109] |
| ***A. paludosum* DSM 8237<sup>T</sup>**<br>(WJBD00000000) | 3,696,449 | 40.1 | isolate | 86 | 99.29 | 2.14 | bog sediment | [105] |
| ***A. tundrae* DSM 9173<sup>T</sup>**<br>(WJBB00000000) | 3,567,344 | 39.7 | isolate | 94 | 99.29 | 1.43 | tundra wetland soil | [110] |
| ***A. wieringae***<br>(LKEU00000000.1) | 3,895,828 | 44.1 | isolate | 62 | 98.57 | 0.71 | sewage digester | [74] |
| ***A. sp. UBA5834***<br>(DJGY00000000.1) | 3,463,311 | 44.1 | MAG | 87 | 98.57 | 1.43 | bioreactor | [111] |
| ***A. fimetarium* DSM 8238<sup>T</sup>**<br>(WJBC00000000) | 3,245,738 | 44.7 | isolate | 99 | 98.57 | 0.71 | cattle manure | [105] |
| ***A. sp. UBA6819***<br>(DKEP00000000.1) | 3,607,361 | 42.8 | MAG | 474 | 94.52 | 3.83 | waste water | [111] |
| ***A. sp. UBA5558***<br>(DIMU00000000.1) | 2,612,337 | 43.6 | MAG | 110 | 85 | 0.71 | bioreactor | [111] |
| ***A. sp. UBA11218***<br>(DNHO00000000.1) | 2,224,242 | 44.8 | MAG | 1,582 | 57 | 0.75 | bioreactor | [111] |

Outside of the type strain, not much is known about the *Acetobacterium* genus. The genus appears to have varied metabolic capabilities, including low-temperature growth [32], debromination of polybrominated diphenyl ethers (PBDEs) [33], hexahydro-1,3,5-trinitro-1,3,5-triazine (RDX) degradation [34, 35], electrode-mediated acetogenesis [36], enhanced iron corrosion [37], and isoprene degradation [38]. To provide a more robust genome dataset, we sequenced and manually curated the genomes of four *Acetobacterium* strains, including *A. malicum* and three psychrophilic strains (*A. fimetarium, A. paludosum,* and *A. tundrae*).

We conducted a comparative genomic study to shed light on the phylogenetic relatedness and functional potential of the *Acetobacterium* genus. Our results suggest that while the major metabolic pathways are well conserved (*e.g.* the Wood-Ljungdahl pathway and accessory components), certain predicted metabolisms and genome features were markedly different than what is known from *Acetobacterium woodii.* For example, a clade consisting of *A.* sp. MES1, *A.* sp. UBA5558, *A.* sp. UBA5834, and *A. wieringae* contained over 40 unique predicted protein sequences with six closely related to diguanylate cyclases, an enzyme that catalyzes the production of the secondary messenger cyclic-di-GMP known to induce biofilm formation [39] and increase tolerance to reactive oxygen species [40]. The unique diguanylate cyclases may afford a unique ability of this clade for cellular attachment to a diverse range of solid surfaces, including carbon-based electrodes. Furthermore, the psychrophilic strains had varied amino acid transport and metabolism, lacking a glycine cleavage system and an alanine degradation pathway, suggesting important protein adaptations to a low-temperature lifestyle. Results described herein provide insight into the defining common features of the *Acetobacterium* genus as well as ancillary pathways that may operate to provide various strains the ability to survive in diverse environments and potentially be exploited for biotechnological applications, such as the production of fuels and chemicals from CO_2_-feedstocks [19, 21].

## Materials and Methods

### DNA extraction, DNA sequencing, and genome assembly of four *Acetobacterium* isolates

For *A. fimetarium, A. malicum, A. paludosum,* and *A. tundrae*, freeze-dried cells were obtained from ATCC and reconstituted on minimal freshwater medium growing autotrophically on H_2_:CO_2_ [12]. Chromosomal DNA was extracted using the AllPrep DNA/RNA Mini Kit (Qiagen) according to manufacturer’s protocol. Extracted DNA was processed for Illumina sequencing using the Nextera XT protocol (Illumina). Samples were barcoded to enable multiplex sequencing of four samples using a single MiSeq V3 kit (2 x 301). Raw paired-end sequences were quality trimmed with CLC Genomics workbench (Qiagen) with a quality score cutoff of Q30, resulting in a total of 297, 911, 471, and 548 million base pairs for *A. fimetarium, A. malicum, A. paludosum,* and *A. tundrae,* respectively. Trimmed paired-end reads were assembled with SPAdes (v. 3.7.0) using the –careful flag to reduce mismatches and short indels [41]. Assembly of trimmed paired-end reads resulted in four high-quality draft genomes, with >90% completion, <5% contamination, the presence of 23S, 16S, and 5S rRNA genes, and at least 18 tRNAs [42] (Table 1; Supplemental Table 1). The SPAdes assembled contigs were assessed for quality using Quast [43] and CheckM (v. 1.0.7) [44].

### Genome completeness

Each *Acetobacterium* genome or metagenome-assembled genome (MAG) was assessed for completeness and contamination (Table 1) using the available *Eubacteriaceae* marker gene set from CheckM (version 1.0.13) [44].

### Manual genome curation and pathway identification

The *Acetobacterium woodii* genome (https://www.ncbi.nlm.nih.gov/nuccore/379009891?report=genbank) was utilized for pathway identification, gene synteny, and sequence similarity. Fasta files of all genomes were downloaded from the NCBI genome database (August, 2018) and uploaded to RAST [45, 46] for annotation. The RASTtk pipeline was used to analyze each genome. The BLAST algorithm in RAST was used to find specific gene sequences and to assess gene synteny of operons (*e.g., rnfCDGEAB*). Sequence similarity of predicted protein sequences are presented in the Supplemental File. Pathways were categorized as highly conserved (high sequence similarity and near identical gene arrangement), conserved (high to medium sequence similarity with variation in gene arrangement), or divergent (low sequence similarity with variation in gene arrangement). To determine what cutoffs should be utilized in assessing pathway conservation, we analyzed RecA across all sequenced *Acetobacterium* strains and found the sequence identity ranged from 100% to 86%. Thus, well-conserved sequences should share at least 86% identity. RAST protein encoding gene (peg) identifiers reveal the synteny of genes that encode each protein in an operon and where available, NCBI accession numbers are provided.

### Concatenated and single protein trees

Predicted protein sequences encoded by the *hydABDEC* operon, RNF operon or individual diguanylate cyclases (DGs) were identified in each genome. Proteins encoded by each operon were manually concatenated into a single contiguous amino acid sequence. Concatenated protein sequences were aligned in MEGA6.06 [47] using MUSCLE [48] with the following parameters: Gap open penalty (−2.9), Gap extend penalty (−0.01), hydrophobicity multiplier (1.2), and UPGMB clustering method with a min. diag. length (lambda) of 24. A protein tree was constructed using the Maximum Likelihood method with the following parameters: test of phylogeny = bootstrap method, number of bootstrap replications = 1000, substitutions type = amino acid, model/method = Jones-Taylor-Thornton (JTT) model, rates among sites = uniform rates, gap/missing data treatment = partial deletion, ML heuristic method = nearest-neighbor-interchange (NNI), and branch swap filter = very strong.

### Pan-genome analysis

Pan-genomics was performed with the Bacterial Pan Genome Analysis Tool (BPGA) version 1.0.0 [49]. Initially, default BPGA parameters were utilized, which includes protein clustering at 50% sequence identity cutoff with USEARCH [50]. Further analysis was performed at various clustering cutoffs ranging from 10% to 99% to examine the pan-genome partitioning (*e.g.,* total gene families, core gene families, accessory gene families, and unique genes). Resulting pan-genome and core genome trees were visualized using FigTree (http://tree.bio.ed.ac.uk/software/figtree/).

The representative *Acetobacterium* core gene sequences and *Eubacteriaceae* core gene sequences from BPGA were uploaded to BlastKOALA [51] to evaluate KEGG pathway predictions. The *Acetobacterium* query dataset contained 1,406 gene families and the *Eubacteriaceae* query dataset contained 44 gene families. Clade-specific gene families were analyzed with BLASTp.

### Data Availability

The sequenced strains were deposited in the sequence read archive (SRA) under the accession numbers WJBE00000000, WJBD00000000, WJBC00000000, and WJBB00000000. KBase narratives for *Acetobacterium* genomes can be found at: https://narrative.kbase.us/narrative/ws.53630.obj.1.

## Results and Discussion

### Phylogeny of the *Acetobacterium* genus

As a first step towards identifying the taxonomic placement of newly sequenced *Acetobacterium* genomes, we collected 16S rRNA gene sequences from the NCBI database, and for MAGs not in the database; we attempted to extract 16S rRNA gene sequences from metagenome data. Multiple MAGs did not contain 16S rRNA gene sequence, potentially due to metagenome assembly and binning issues. *A. dehalogenans* contained a truncated 16S rRNA gene that was less than 400 bp. Therefore, nine 16S rRNA gene sequences of the 13 sequenced *Acetobacterium* genomes were utilized. We reconstructed an updated phylogenetic tree of available 16S rRNA gene sequences (Figure 1). The *Acetobacterium* genus forms distinct subgroup with cluster XV of the *Clostridium* subphylum [52]. Notably, Willems and Collins found that the psychrophilic strains were not more closely related to one another compared to non-psychrophilic strains—our findings suggest the opposite, with distinct clustering of known psychrophiles *A. tundrae, A. paludosum, A. fimetarium, and A. bakii* (Figure 1). Other *Acetobacterium* strains with sequenced genomes clustered close together, including *A. wieringae* and *A.* sp. MES1.

**Figure 1.**
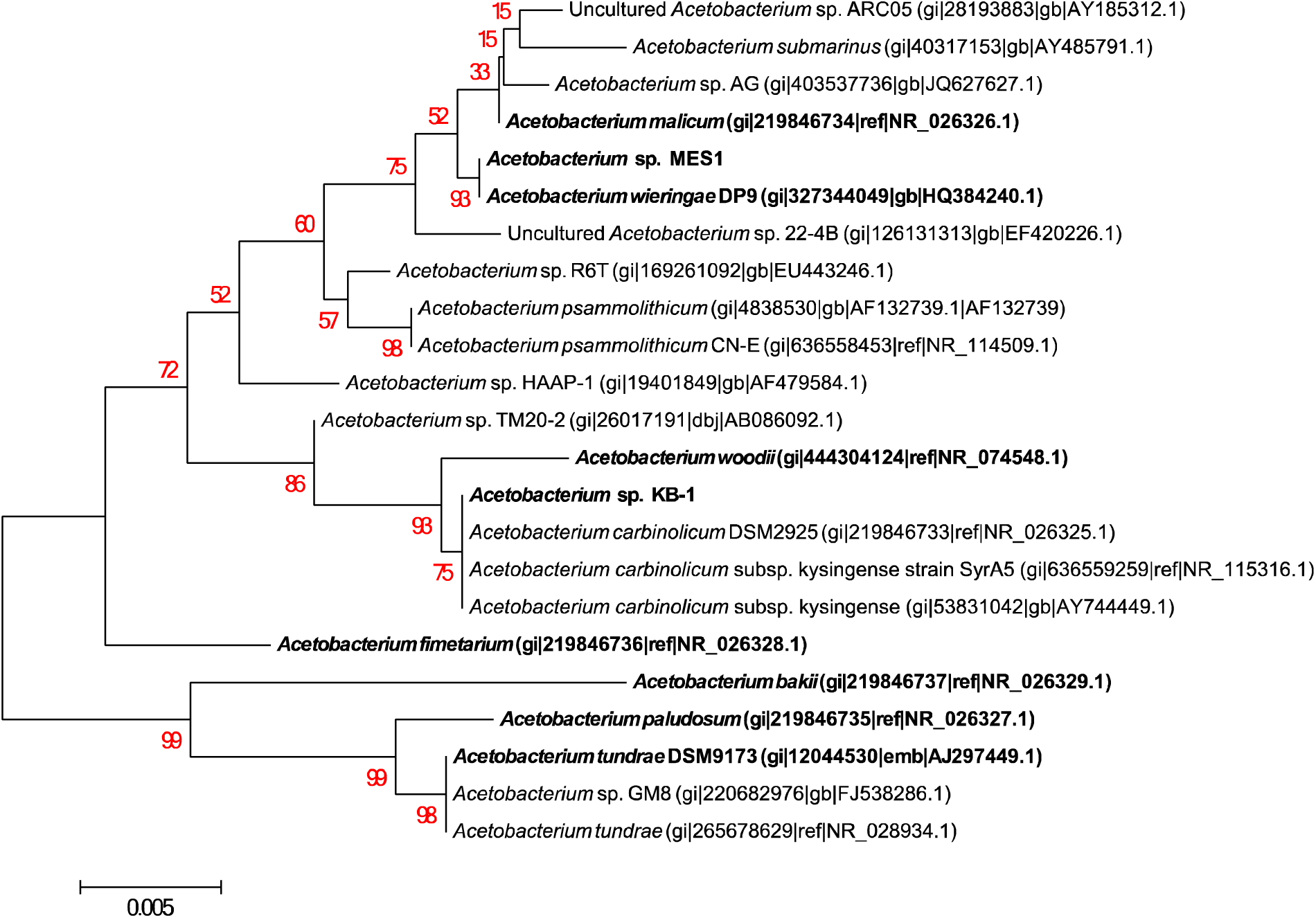
16S rRNA gene phylogeny. Phylogenetic tree of *Acetobacterium* 16S rRNA gene sequences. *Acetobacterium* strains in bold represent strains that have a sequenced genome. Numbers in red indicate percent bootstrap support.

### Genome characteristics of the *Acetobacterium* genus

There are 14 sequenced representatives of the *Acetobacterium* genus. Thirteen of the 14 sequenced *Acetobacterium* strains are draft genome assemblies, with the genome of *Acetobacterium woodii* as the only complete and closed genome (contained in one contiguous sequence). The thirteen draft genomes had an average GC percent of 42.7 ± 1.9% and were comprised of a varying number of contigs, ranging from 62 (*A. wieringae*) to 1,006 (*A.* sp. KB-1) and the genome sizes ranged from ∼3.2 Mbp (*A. fimetarium*) to ∼4.1 Mbp (*A. bakii*) (Table 1). Each genome was examined for completeness based upon the *Eubacteriaceae* marker gene set from CheckM (Parks et al., 2015). The fourteen *Acetobacterium* genomes/MAGs were >98% complete with the exception of *A.* sp. UBA6819 (94% complete, missing 11 marker genes), *A.* sp. UBA5558 (85% complete, missing 31 marker genes), and *A.* sp. UBA11218 (57% complete, missing 93 marker genes). *A.* sp. UBA5558 was only 2.61 Mbp, well below the average genome size (3.69 Mbp), and a genome coverage of 13x. Likewise, *A.* sp. UBA11218 was only 2.22 Mbp with 6.1x genome coverage. These genomes were often excluded from our analyses in order to more accurately describe the core components of the genus.

To understand the relatedness of the *Acetobacterium* genus, we calculated the pairwise average nucleotide identity (ANI) and average amino acid identity (AAI) for each genome and constructed phylogenies based on whole genome alignments (Figure 2). The closest species to *Acetobacterium woodii,* the type strain of the genus, were *A. malicum, A. dehalogenans, A.* sp. KB-1, with average nucleotide identity (ANI) and average amino acid identity (AAI) of 80% and 79%, respectively. In contrast, many of the other *Acetobacterium* genomes were highly similar to one another, potentially representing subspecies in the genus. Specifically, *A.* sp. UBA6819 and *A.* sp. KB-1 (99% ANI); *A. malicum,* and *A. dehalogenans* (97% ANI); *A.* sp. MES1, *A.* wieringae, *A.* sp. UBA5558, and *A.* sp. UBA5834 (97-100% ANI); and *A. tundrae* and *A. paludosum* (95% ANI) had high sequence similarity across their genomes (Figure 2; [53]). These findings suggest that, for example, three potential subspecies groups exist: 1) *A. malicum* and *A. dehalogenans,* 2) *A*. sp. UBA6819 and *A.* sp. KB-1, and 3) *A.* wieringae, *A.* sp. MES1, *A.* sp. UBA5558, and *A.* sp. UBA5834 (brackets in Figure 2 indicate suggested strain-level variation of a single species). While many of the MAGs were above the suggested species-level cutoff for ANI of 95% [54–56], we cannot definitively define particular MAGs as subspecies and phenotypic characterization should accompany genomic data to confirm these findings.

**Figure 2.**
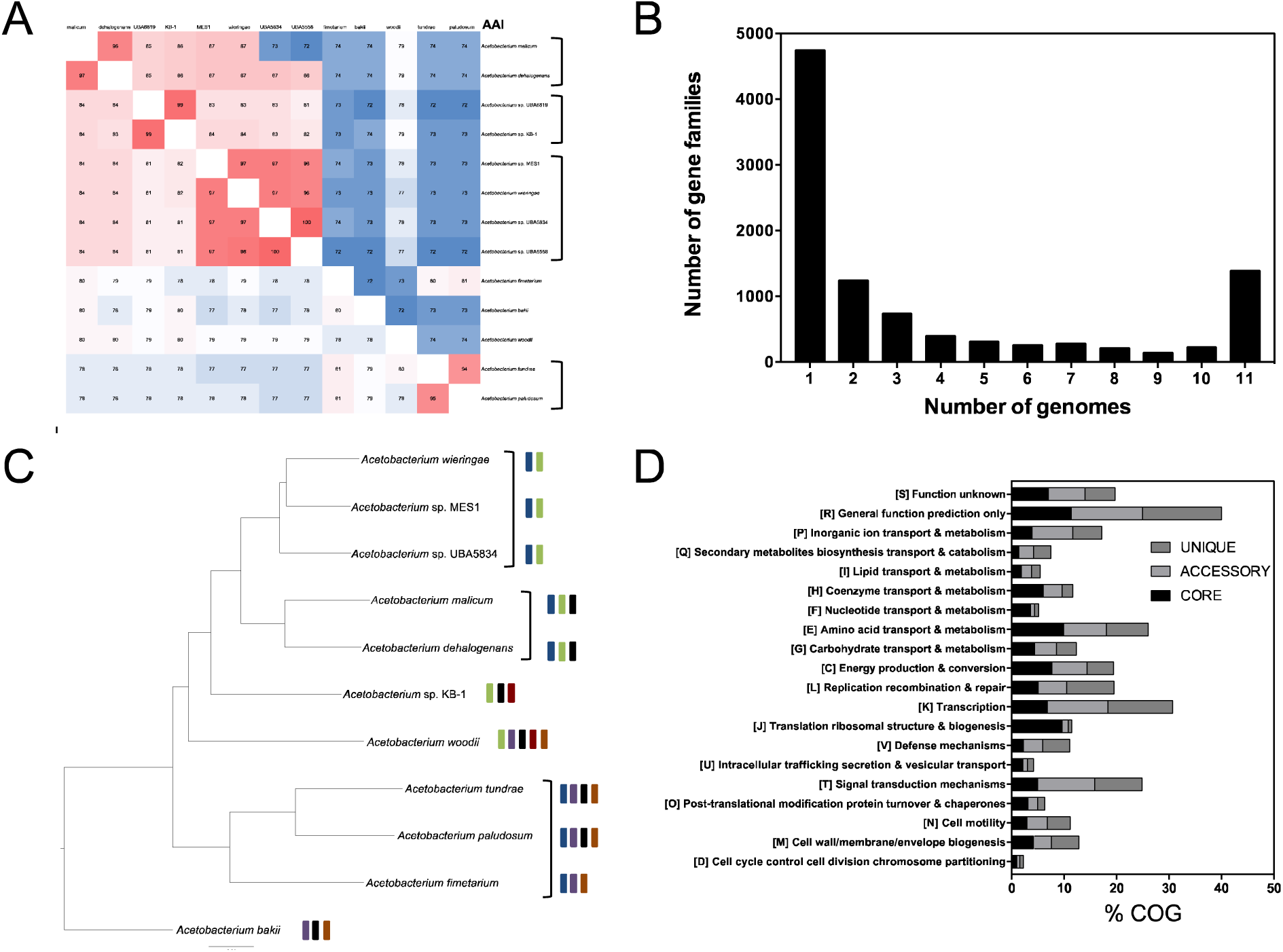
*Acetobacterium* pan-genome. A) ANI/AAI matrix of sequenced *Acetobacterium* isolate genomes and MAGs. Potential sub-species are denoted in brackets. B) Pan-genome distribution of gene families found in 1, some (2-10), or all genomes (determined using predicted amino acid sequences). C) Pan-genome phylogeny of 11 *Acetobacterium* strains. Potential sub-species denoted in brackets. Blue bar = presence of Rnf2 complex, green bar = potential for 2,3-butanediol metabolism, purple bar = *A. woodii-*type WLP cluster I, black bar = potential for methanol oxidation, orange bar = potential for caffeate metabolism, red bar = potential for ethanol oxidation D) COG classification of predicted protein sequences of the core genome, accessory genome, and unique genome of 11 *Acetobacterium* strains.

### The *Acetobacterium* pan-genome

Utilizing predicted protein sequences, whole genome comparisons between *Acetobacterium* genomes was performed. While the presence of a particular pathway does not definitively prescribe function, genetic comparisons reveal evolutionary conservation or divergence and provide valuable information about the functional potential encoded in each genome. Pan-genome analyses utilized gene families or amino acid sequences that clustered together at a defined sequence similarity cutoff. We identified the common set of gene families (core genome), the unique set of gene families (unique genome), and the ancillary set of gene families, which were found in at least two, but not all genomes (accessory genome). In order to obtain the most accurate pan-genome and core genome estimates, only genomes with a >98% completion were utilized (11 in total).

Whole genome comparisons corroborated 16S rRNA gene phylogeny and ANI/AAI groupings of the *Acetobacterium* genus (Figure 2). Inclusion of the eleven most complete *Acetobacterium* genomes revealed a total of 10,126 gene families (40,203 total amino acid sequences). When calculated as the total number of amino acid sequences per category, the percentage of core sequences was 38.4%, while the accessory, unique, and exclusively absent sequences were 43.6%, 11.9%, and 6.1%, respectively. Within the pan-genome, the core genome, accessory genome, and unique genome contained a total of 1406 (13.9%), 3,964 (39.1%), and 4,756 (46.9%) gene families, respectively (Figure 1B, Table 2). The pan-genome partitioning of gene families suggests that the unique genome contains the most functional diversity, followed by the accessory genome and then the core genome. Phylogenomic trees utilizing the pan-genome and core genome sequences revealed clustering of the psychrophilic strains *A. tundrae, A. paludosum,* and *A. fimetarium* (Figure 1C). Other clusters included *Acetobacterium wieringae, A.* sp. MES1, and *A.* sp. UBA5834; and *A. malicum* and *A. dehalogenans*, consistent with the ANI and AAI results.

**Table 2.** Pan-genome distribution of the eleven most complete *Acetobacterium* genomes.

|  | **Core gene families** | **Accessory gene families** | **Unique gene families** | **Exclusively absent gene families\*** |
| --- | --- | --- | --- | --- |
| *A. bakii* | 1406 | 1577 | 811 | 57 |
| *A. dehalogenans* | 1406 | 1814 | 358 | 0 |
| *A. fimetarium* | 1406 | 1070 | 510 | 83 |
| *A. sp. KB-1* | 1406 | 1631 | 448 | 17 |
| *A. malicum* | 1406 | 1915 | 342 | 3 |
| *A. sp. MES1* | 1406 | 1680 | 223 | 3 |
| *A. paludosum* | 1406 | 1497 | 483 | 1 |
| *A. tundrae* | 1406 | 1421 | 486 | 28 |
| *A. sp. UBA5834* | 1406 | 1505 | 222 | 26 |
| *A. wieringae* | 1406 | 1860 | 253 | 0 |
| *A. woodii* | 1406 | 1548 | 620 | 27 |

\*Exclusively absent gene families are those found in the denoted strain only.

Protein sequences from core, accessory, and unique gene families were classified with the clusters of orthologous groups (COG) database [57]. The core genome had the highest representation from categories E (amino acid transport & metabolism), J (translation, ribosomal structure & biogenesis), and C (energy production and conversion) (Figure 1D). The highest representation in the accessory and unique gene families were from Category K (transcription) and category T (signal transduction mechanisms). The third most represented category for the accessory and unique gene families were category E (amino acid transport and metabolism) and category L (replication, recombination, and repair), respectively. Sequences were also characterized with the KEGG database (Supplemental File).

### The Wood-Ljungdahl (acetyl-CoA) carbon fixation pathway in Acetobacterium spp

The Wood-Ljungdahl pathway (WLP) (or the reductive acetyl-CoA pathway) is thought to be derived from an ancient carbon dioxide fixing metabolic pathway [58]. The WLP is coupled to substrate-level phosphorylation but the net ATP gain is zero, thus a chemiosmotic mechanism is employed for energy conservation [59]. Generation of a transmembrane gradient is utilized by an F_1_F_0_ ATP synthase for ATP synthesis and relies upon either the *Rhodobacter* nitrogen fixation (Rnf) complex or cytochromes depending upon the species. The *Acetobacterium* genus appears to lack cytochromes and therefore *Acetobacterium woodii* has served as a model organism for understanding the mechanisms of energy conservation in acetogens lacking cytochromes [22, 60–64].

#### Cluster I of the WLP is divergent across *Acetobacterium* spp

In *Acetobacterium woodii,* the WLP is organized into three separate and distinct gene clusters [22]. Cluster I contains genes that encode the <u>h</u>ydrogen <u>d</u>ependent <u>C</u>O_2_ reductase (HDCR), which is the first step of the WLP methyl branch (CO_2_ to formate) [65]. Specifically, Cluster I in *Acetobacterium woodii* encodes for formate dehydrogenase (FDH), formate dehydrogenase accessory protein, a putative FeS-containing electron transfer protein, and an [FeFe]-hydrogenase (FdhF1, HycB1, FdhF2, HycB2, FdhD, HycB3, and HydA2) [22].

The presence and arrangement of Cluster I was determined for the remaining 12 MAGs, and this cluster was only present in *A. bakii, A. fimetarium, A. paludosum,* and *A. tundrae* (Figure 3; Supplemental File), findings consistent with recent work by Esposito and coworkers [53]. Furthermore, *A. fimetarium* only encodes for FdhF1 and not FdhF2. Cluster I was markedly different and more divergent in the remaining genomes, with low sequence identity (<50%) and similarity (Supplemental Figure 1 and Supplemental file). Phylogenetic analysis of FdhF1 from *A. woodii* revealed clustering with FdhF1 from *A. bakii, A. paludosum,* and *A. tundrae*, and formate dehydrogenases from two strains from the Lachnospiracaea family, *Blautia schinkii* and *Blautia hansenii* (71% identity, 98% coverage) (Supplemental Figure 1). The remaining *Acetobacterium* strains encode for an NAD-dependent formate dehydrogenase, which clustered together and with *Eubacterium limosum* (76% identity; 99% coverage) or *Alkalibaculum bacchi,* both from the *Eubacteriaceae* family.

**Figure 3.**
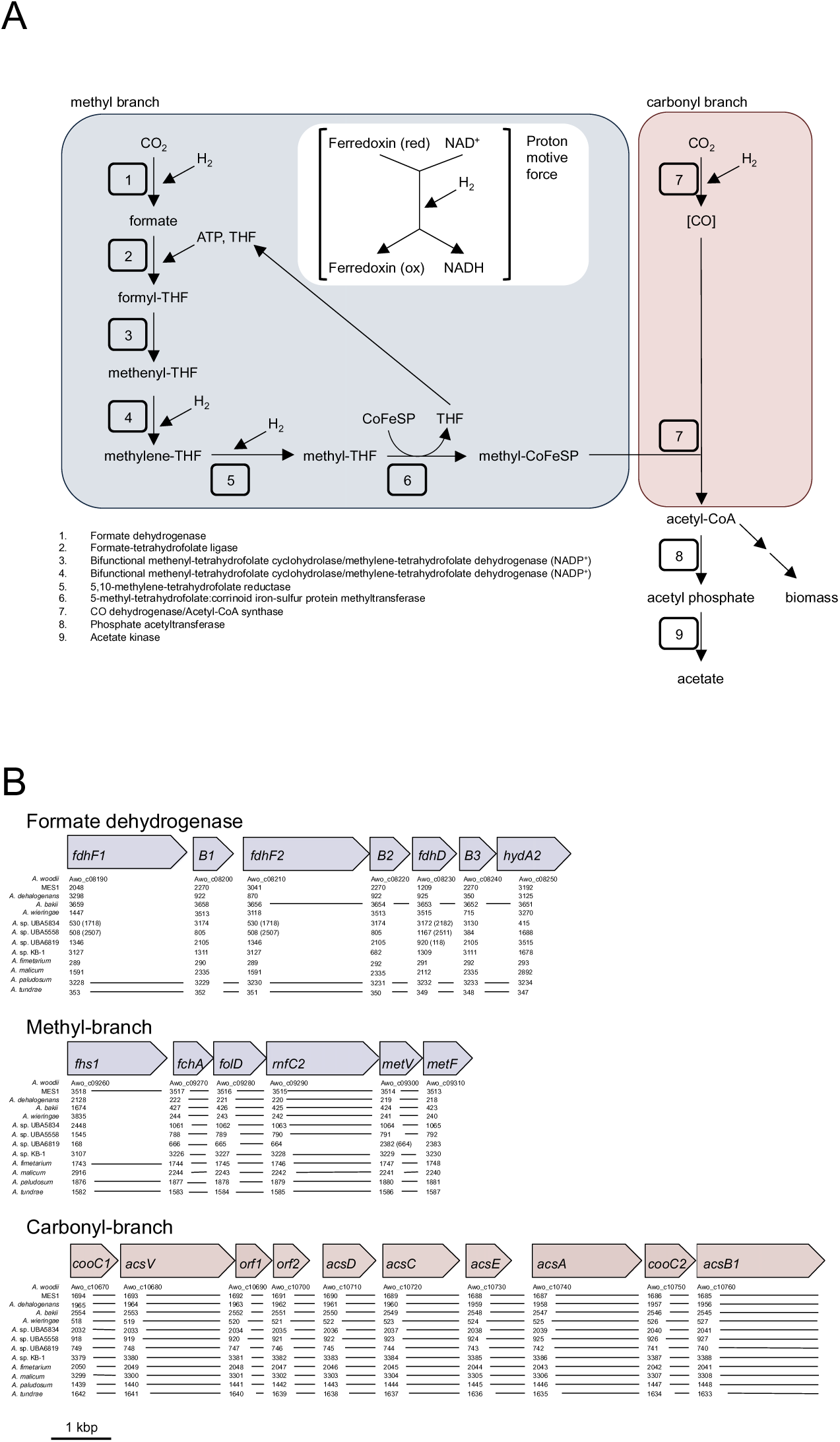
The Wood-Ljungdahl pathway (WLP). A) Pathway diagram of the methyl and carbonyl branches of the WLP. B) Genetic representation of the WLP cluster I (formate dehydrogenase), Cluster II (Methyl branch), and Cluster III (Carbonyl branch). Lines denote contiguous sequences, while the absence of a line represents a discontiguous operon.

The variability in FDH could be a result of the availability and requirement for tungsten. In *Campylobacter jejuni*, and *Eubacterium acidaminophilum*, formate dehydrogenase activity is dependent upon tungsten availability, as *tupA* mutants exhibited markedly decreased FDH activity [66, 67]. We found that the strains encoding for the NAD-dependent formate dehydrogenase also encode for a tungsten-specific ABC transporter (*tupABC*). TupA has a high affinity for tungsten and can selectively bind tungsten over molybdate [68]. The *tupABC* operon was absent in the strains encoding the *A. woodii*-type Cluster I, though the ability to transport tungsten in these strains may rely upon the *modABC* operon, as all sequenced genomes encode for ModA, which is a high affinity molybdate ABC transporter that binds both molybdate and tungstate [66, 69].

#### Conservation of WLP Cluster II and Cluster III of the *Acetobacterium* genus

Cluster II (Methyl branch) encodes for enzymes responsible for the conversion of formate to methyltetrahydrofolate (methyl-THF) (Fhs1, FchA, FolD, RnfC2, MetV, and MetF). Unlike Cluster I, Cluster II is highly conserved across all *Acetobacterium* genomes (Supplemental File). The exception was the dihydrolipoamide dehydrogenase (LpdA1) and glycine cleavage system H protein (GcvH1), which is found downstream of methylene-tetrahydrofolate reductase (MetF) in *A. woodii* but dispersed throughout the genomes of the other MAGs (Figure 3; Supplemental File). Other variations in Cluster II are found in *A. tundrae* and *A. paludosum,* which contain two formate tetrahydrofolate-ligase (Fhs1) subunits separated by a small hypothetical protein (39 aa). In *A. paludosum* the two copies are identical, while in *A. tundrae* there is one amino acid difference between the two Fhs1 sequences, suggesting a gene duplication event in these strains.

Cluster III (Carbonyl branch) encodes for the CO dehydrogenase/acetyl-CoA synthase complex (AcsABCD), methyltransferase (AcsE), and accessory proteins (CooC and AcsV), for conversion of methyl-THF to acetyl-CoA. The Carbonyl branch is the most well conserved portion of the WLP in the *Acetobacterium* genus, with high sequence similarity and identical gene arrangement across all genomes—the one exception is a hypothetical protein and an additional AcsB directly downstream of AcsB in *A. bakii, A. dehalogenans, A. paludosum, A. tundrae, A. fimetarium, A. malicum,* and *A.* sp. KB-1 (Figure 2; [32]). The CO dehydrogenase/acetyl-CoA synthase complex (AcsABCD) converts CO_2_ to CO and this conversion has the largest thermodynamic barrier of the WLP. Acetogens utilize flavin-based electron bifurcation to overcome this barrier [1]. This highly specialized form of energy conservation may explain why this cluster is well conserved, demonstrating that purifying selection is the predominant evolutionary force on this cluster.

### Genes essential for energy conservation

Energy conservation in acetogens requires the generation of a sodium (cytochrome-deficient acetogens like *Acetobacterium*) or proton gradient (cytochrome-containing acetogens) across the cytoplasmic membrane [1]. In *Acetobacterium woodii*, and potentially all other members of the genus, a sodium-ion gradient is utilized. The transmembrane sodium-ion gradient is generated by the Rnf complex, but a number of accompanying enzyme complexes, including an electron bifurcating hydrogenase, ferredoxins, and electron-transfer flavoproteins (ETF) are also essential for overcoming thermodynamically unfavorable reactions of the WLP and for ATP generation [1].

#### Conservation of the flavin-based electron bifurcating hydrogenase HydABC

The key hydrogenase responsible for providing reducing equivalents for energy conservation in *Acetobacterium woodii* is encoded by *hydA1*, *hydB*, *hydD*, *hydE*, and *hydC* (Awo_c26970-Awo_c27010). This electron-bifurcating hydrogenase, HydABCDE, couples the thermodynamically unfavorable reduction of oxidized ferredoxin with the favorable reduction of NAD^+^ [22]. With the exception of *A.* sp. UBA5558, HydABCDE (Supplemental File) was well conserved across the *Acetobacterium* genus (>77% identity and 85% coverage). A phylogenetic tree of concatenated proteins HydABCDE revealed identical clustering to that of the whole genome tree, with tight clustering of *A.* sp. MES, *A.* sp. UBA5834, and *A. wieringae*, and clustering of the psychrophilic strains (Supplemental Figure 2). This high degree of conservation across the genus highlights the essential function of this enzyme complex in *Acetobacterium* spp.

#### Conservation of the ferredoxin-NAD^+^ oxidoreductase (Rnf complex) and variability in Rnf copy number

Energy conservation in *Acetobacterium woodii* requires a multi-subunit integral membrane ferredoxin-NAD^+^ oxidoreductase called the *<u>R</u>hodobacter* <u>n</u>itrogen fixation (Rnf) complex [70–72]. The Rnf complex is an energy-coupled transhydrogenase responsible for transfer of electrons from reduced ferredoxin to NAD^+^, which generates a transmembrane sodium ion gradient to drive ATP generation via sodium-dependent ATP synthase. This process is reversible when the concentration of NADH is greater than ferredoxin [64]. The Rnf cluster is common across many *Clostridia* including *C. tetani, C. kluyveri, C. difficile, C. phytofermentans, and C. botulinum* [60], however the *Clostridia*-type is not sodium-dependent and may involve a transmembrane proton gradient [59]. While gene arrangement within the Rnf cluster differs across prokaryotes, in *Acetobacterium woodii* the Rnf genes (*rnfCDGEAB*) are polycistronic and co-transcribed [60]. All sequenced *Acetobacterium* genomes encode for a Rnf complex (Supplemental File). Based upon the presence and conservation of the Rnf complex in sequenced *Acetobacterium* spp., we propose that energy conservation mechanisms are similar across the genus and that Rnf is required.

Microorganisms, such as *Azotobacter vinelandii*, encode for two Rnf complexes, one of which is linked to nitrogen fixation and the other that is expressed independent of nitrogen source [73]. *Acetobacterium woodii* only encodes for a single Rnf complex [22], while *Acetobacterium wieringae* and *Acetobacterium* sp. MES1 were found to encode for two [14, 74]. The function of the second Rnf complex in *Acetobacterium* is unknown, but closer examination of the remaining *Acetobacterium* genomes revealed that the majority of *Acetobacterium* genomes encode for a second *rnfCDGEAB* cluster (Supplemental File). The genome architecture surrounding the second Rnf complex was also well conserved (Supplemental data), with a predicted hydroxymethylpyrimidine transporter, hydroxyethylthiazole kinase, and thiamin-phosphate pyrophosphorylase directly adjacent to the Rnf2 complex, suggesting a potential role in thiamin pyrophosphate (Vitamin B_1_) synthesis and nitrogen metabolism [75]. The genomes encoding only the first Rnf complex retained the genes that surround the second Rnf complex, supporting an HGT event. Phylogenetic analysis supports these findings, as the second Rnf cluster forms a distinct clade with the *Clostridia* Rnf cluster (Figure 4). Therefore, the first Rnf cluster is most likely a paralog from a *Eubacteriaceae* ancestor, while the second copy is a potential ortholog from a horizontal gene transfer (HGT) event.

**Figure 4.**
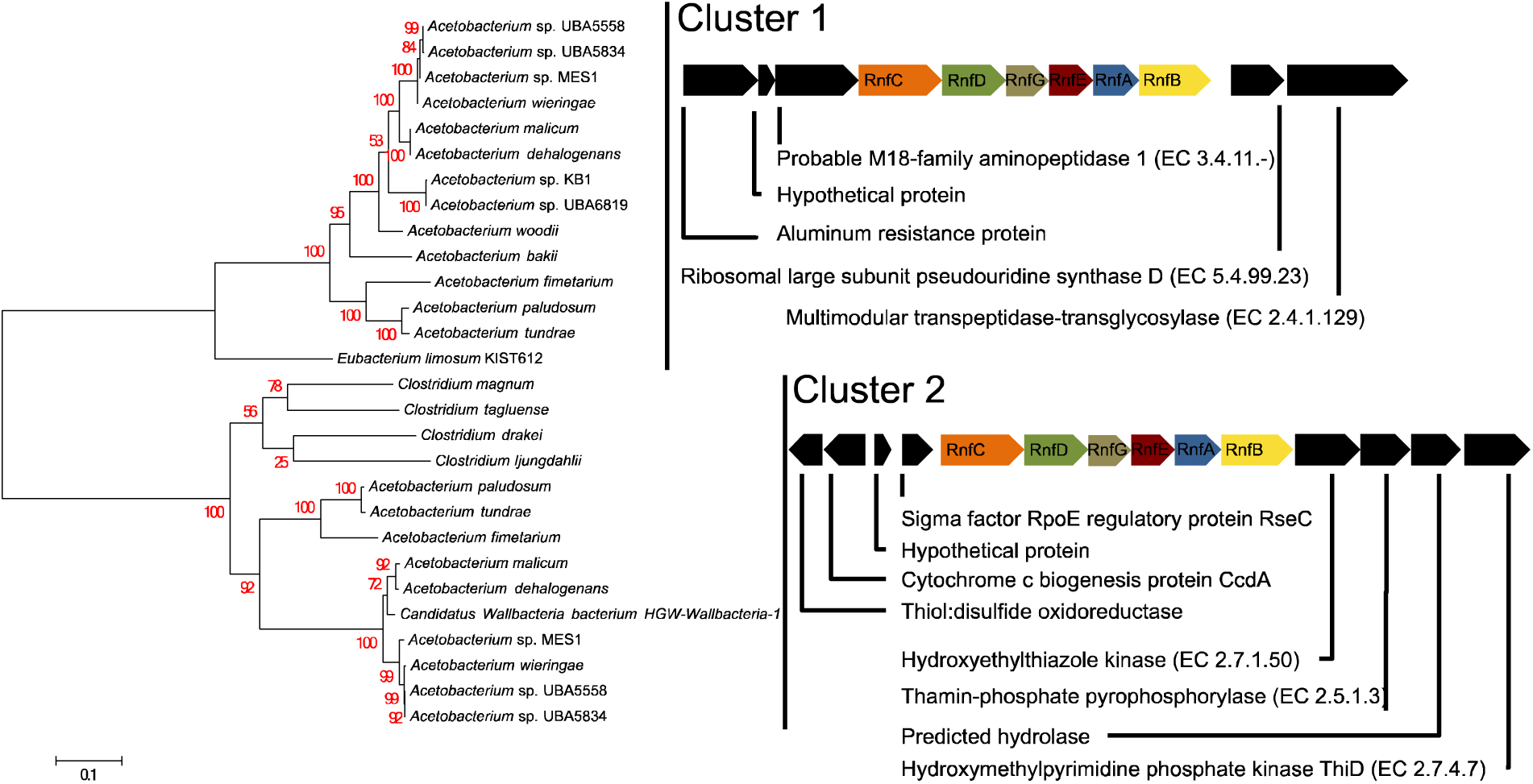
Phylogenetic tree of the RnfCDGEAB protein complex. Predicted protein sequences were concatenated and clustered with MUSCLE [48] and the phylogenomic tree was constructed with the Maximum Likelihood method using the Bootstrap method with 1000 replications for test of phylogeny. The substitution method was Jones-Taylor-Thornton with uniform rates among sites. Clustering and tree construction was performed with MEGA 6.06 [47]. Varied genome architecture of predicted protein sequences immediately surrounding each Rnf operon are shown (not drawn to scale).

#### Conservation of the Sodium-dependent ATPase

*Acetobacterium woodii* encodes for an integral membrane Na^+^ F_1_F_0_ ATP synthase that generates ATP via the sodium ion gradient and contains both V-type and F-type rotor subunits [76]. All sequenced *Acetobacterium* genomes encode for the F-type and V-type ATP synthase (Supplemental File), each contained in a separate operon and with amino acid sequence similarity to *Acetobacterium woodii* ranging from 56% (AtpB, *A. fimetarium*) to 98% (AtpE2, *A. bakii*). In particular, the *c*-subunits of the F_1_F_0_ ATP synthase are responsible for ion translocation across the membrane. Using three *c-* subunits from *Acetobacterium woodii* (Awo_c02160 – c02180), which is known to use a sodium ion gradient, we queried the remaining 12 *Acetobacterium* genomes for *c*-subunits, and specifically, the Na^+^-binding motif. All strains encode for at least two subunits with binding motifs specific for Na^+^ (Supplemental Figure 3). The presence of the F-type and V-type ATP synthase operons, gene synteny, and sequence similarity across all strains, in addition to the presence of Na^+^-binding motifs suggests that all *Acetobacterium* strains utilize this ATP synthase for ATP production via a sodium gradient, similar to what is observed in *A. woodii*.

### Other genome features

#### Alternative Electron Donors/Acceptors utilized by the *Acetobacterium* genus

*Acetobacterium* species can use a wide range of substrates for carbon and energy, indicating a generalist lifestyle for the genus despite the perception that acetogens are specialists. This could provide an advantage in anoxic environments where competition for limited substrates is high [77, 78]. As an example, encoded in the genomes of most *Acetobacterium* species are carbon utilization pathways for caffeate, 1,2-propanediol, 2,3-butanediol, ethanol, lactate, alanine, methanol, glucose, fructose, and glycine betaine (Supplementary File).

Acetogens metabolize alcohols as an alternative to autotrophic acetogenic growth [79]. *Acetobacterium woodii* can utilize 1,2-propanediol as the sole carbon and energy source for growth [30]. In anaerobic environments, formation of 1,2-propanediol results from the degradation of fucose and rhamnose, constituents of bacterial exopolysaccharides and plant cell walls [80]. The 1,2-propanediol degradation pathway is encoded by the *pduABCDEGHKL* gene cluster, which in *A. woodii* (Awo_c25930-Awo_c25740) contains 20 genes with similarity to the *pdu* cluster of *Salmonella enterica* [30]. The presence, homology, and gene arrangement of the *pdu* gene cluster in each of the other 12 *Acetobacterium* genomes suggests that 1,2-propanediol degradation is conserved across the *Acetobacterium* genus (Supplemental File). Furthermore, all strains contain a histidine kinase and response regulator upstream of the 1,2-propanediol cluster suggesting that the regulatory and expression mechanisms of this pathway are conserved. In *A. woodii*, it was proposed to sense alcohols (chain length > 2) or aldehyde intermediates—a mechanism we hypothesize is employed by all *Acetobacterium* species.

The 2,3-butanediol oxidation pathway is encoded by the *acoRABCL* operon in *Acetobacterium woodii* and unlike 1,2-propanediol degradation, the WLP accepts reducing equivalents from 2,3-butanediol oxidation to generate acetate [29]. AcoR is a putative transcriptional activator, AcoA is a TPP-dependent acetoin dehydrogenase (alpha subunit), AcoB is a TPP-dependent acetoin dehydrogenase (beta subunit), AcoC is a dihydrolipoamide acetyltransferase, and AcoL is a dihydrolipoamide dehydrogenase [29, 81, 82]. Amino acid sequence homology and operon structure suggests that 2,3-butanediol oxidation is a common metabolism across the *Acetobacterium* genus, with the exception of *A. fimetarium, A. paludosum,* and *A. tundrae* (Supplemental File). *A. bakii* encodes for this cluster but low percent identity (<50%) and positives (<66%) for the AcoR subunit, AcoC subunit and AcoL subunit. The genome architecture surrounding the 2,3-butanediol operon showed slight variations in some strains, as *A. bakii* has an additional 2,3-butanediol dehydrogenase (WP_050738536.1) and a small hypothetical protein (WP_050738535.1) between AcoR and AcoA (Supplemental File). Likewise, *A. malicum*, *A. dehalogenans*, *A*. sp. MES1, *A. wieringae*, *A.* sp. UBA5558, and *A.* sp. UBA5834 encode for a small hypothetical protein between *acoR* and *acoA*.

In acetogens, the oxidation of primary aliphatic alcohols, such as ethanol, is coupled to the reduction of CO_2_ [83]. The ethanol oxidation pathway has been elucidated in *A. woodii* and contains a bifunctional acetaldehyde-CoA/alcohol dehydrogenase (AdhE) and 7 iron type alcohol dehydrogenases (Adh1-7) (Supplemental File). Interestingly, in one study, only *adhE* from this operon in *A*. *woodii* was upregulated during growth on ethanol [26], suggesting this was the primary enzyme responsible for ethanol oxidation. However, *Acetobacterium* sp. KB-1 and *A.* sp. UBA6819 were the only strains to encode for a complete AdhE with 87% sequence identity to *A. woodii* (Supplemental File). Despite the lack of AdhE in the other species of *Acetobacterium*, species like *A. wieringae* are capable of growth on ethanol [79, 84–86]. It is likely that *A. wieringae* and the other *Acetobacterium* strains that do not encode for the *A. woodii-*type bifunctional acetaldehyde-CoA/alcohol dehydrogenase employ an alternative pathway for ethanol oxidation. Genomic and biochemical analyses are needed to confirm this phenotype but one hypothesis is the conversion of ethanol to acetate proceeds via acetaldehyde by the activities of alcohol dehydrogenase (converts ethanol to acetaldehyde) and aldehyde:ferredoxin oxidoreductase (converts acetaldehyde directly to acetate). This has been shown in *Thermacetogenium phaeum* [87]. The genome of *A.* sp. MES1 encodes for an alcohol dehydrogenase (peg.3269) upstream of a tungsten-containing aldehyde:ferredoxin oxidoreductase (AFO) (peg.3267). *A. woodii* does encode for a similar alcohol dehydrogenase [Adh3; alcohol dehydrogenase, iron-type (AFA47416)], but does not contain the tungsten-containing AFO.

Utilization of lactate for carbon and energy has been observed in many *Acetobacterium* strains, including *A. woodii, A. wieringae, A. dehalogenans, A. carbinolicum, A. malicum, A. fimetarium, A. bakii, A. paludosum,* and *A. tundrae*. The protein machinery responsible for lactate metabolism is encoded by the lctABCDEF operon [88, 89]. The operon includes a transcriptional regulator (LctA), an electron transfer flavoprotein beta (LctB) and alpha (LctC), lactate dehydrogenase (LctD), a potential L-lactate permease (LctE), and a potential lactate racemase (LctF). With the exception of *A.* sp. UBA5558, all sequenced *Acetobacterium* strains encode for the lactate operon (Supplemental File).

Recently, Donig and Muller demonstrated that *A. woodii* is capable of utilizing alanine as a sole carbon and energy source, and identified the pathway responsible for this metabolism [27]. The alanine degradation pathway consists of a pyruvate:ferredoxin (flavodoxin) oxidoreductase (PFO) (AWO_RS12520), a sodium:alanine symporter family protein (AWO_RS12525), alanine dehydrogenase (AWO_RS12530), and Lrp/AsaC family transcriptional regulator (AWO_RS12535). Examination of this operon revealed it was well conserved across the *Acetobacterium* genus, with the exception of *A. fimetarium, A. paludosum,* and *A. tundrae*, which lacked a sodium:alanine symporter, alanine dehydrogenase, and the transcriptional regulator, the details of which are discussed below (Supplemental File).

Some acetogenic bacteria utilize phenyl acrylates as alternative electron acceptors [90]. One such phenyl acrylate, caffeate, is produced during lignin degradation and may be an available substrate in environments containing vegetation [91]. Recently it was observed that *A. woodii* has the ability to couple caffeate reduction with ATP synthesis [28]. The caffeate reduction operon in *A. woodii* is encoded by *carA2*, *carB2*, *carC, carD,* and *carE* (Awo_c15700-Awo_c15740). Specifically, CarA is a hydrocaffeyl-CoA:caffeate CoA transferase [28] CarB is an ATP-dependent acyl-CoA synthetase [92], CarC is a caffeyl-CoA reductase and CarDE is an electron transfer protein [62]. Examination of the other twelve *Acetobacterium* genomes revealed BLAST hits with low similarity (<53% identity) with the exception of three psychrophilic strains (*A. paludosum, A. fimetarium,* and *A. tundrae*) and CarC, CarD, and CarE from *Acetobacterium bakii* (74%, 73%, and 73% amino acid sequence identity; 88%, 85%, and 86% amino acid sequence coverage) (Supplemental File). The lack of similar and congruous sequences suggests only *A. woodii, A. fimetarium, A. paludosum,* and *A. tundrae* are capable of caffeate reduction via this pathway.

#### Potential adaptations for enhanced surface colonization

Many of the *Acetobacterium* genomes/MAGs encoded for portions of the Widespread Colonization Island (WCI), which mediates non-specific adherence to surfaces and biofilm formation [93]. The products of the WCI are responsible for assembly and secretion of bundled pili [94] and may be important for colonization of diverse environments [95]. The general structure of the WCI genomic region is similar for all sequenced *Acetobacterium* strains with the exception of *A. woodii*, and includes 1 to 4 small hypothetical proteins (∼56 a.a), followed by TadZ, Von Willebrand factor type A, RcpC/CpaB, TadZ/CpaE, TadA/VirB11/CpaF, TadB, and TadC (Supplemental File). The small hypothetical proteins from *A.* sp. MES1 show significant similarity (100% coverage, 98% identity) with Flp/Fap pilin component from *A. wieringae* (OFV69504.1-OFV69507.1). Likewise, immediately upstream of the Flp/Fap pilin components is a hypothetical protein with high similarity (100% coverage, 79% identity) to a prepilin peptidase from *A. dehalogenans* (WP_026393143.1). The presence of this potentially biofilm-enhancing WCI found in many of the species of *Acetobacterium* but not in *A. woodii* highlights the importance of expanding genome-informed biochemical analyses to species and strains beyond the type strain of a genus. One intriguing hypothesis resulting from the discovery of this WCI is that it may enhance surface attachment on insoluble electron donors like metallic iron or electrodes. A recent study suggests *A. malicum* and an isolate most closely related to *A. wieringae* more effectively extract electrons from solid Fe(0) coupons compared to *A. woodii* [37]. Additionally, *A*. sp. MES1 is capable of colonizing cathodes in microbial electrosynthesis systems [14, 36], but *A. woodii* has failed in all such attempts [96, 97].

Closer examination of the strains that were phylogenetically most-closely related to *Acetobacterium wieringae*, which includes *A.* sp. MES1 (capable of electroacetogenesis), revealed 42 proteins unique to this clade (Supplemental File). Of these unique protein sequences, several were of potential relevance for possible surface colonization and electron transport, including cardiolipin synthase for the production of membrane phospholipids, the global regulator diguanylate cyclase (Supplemental Figure 7), and methylenetetrahydrofolate reductase (methylene-THF reductase) (Supplemental File).

#### Potential adaptations to a psychrophilic lifestyle

Psychrophilic microorganisms have adapted to survive and grow in cold environments by varying membrane fluidity, optimization of transcription and translation (*e.g.,* overexpression of RNA helicases, post-transcriptional regulation of RNA), and expressing cold shock proteins and cold-adapted enzymes [98]. *Acetobacterium bakii* employs post-transcriptional regulation for cold-adaptation and at low temperatures a lipid biosynthesis pathway (ABAKI_c35860-c35970), cold shock protein CspL (ABAKI_c09820 and ABAKI_c26430), and a Dead-box helicase (ABAKI_c00160-c00180) were upregulated [32]. Examination of these pathways across the *Acetobacterium* genus revealed the lipid biosynthesis pathway was conserved, but for the psychrophilic strains the genome architecture surrounding this cluster was different (Supplemental File). Furthermore, two of the four psychrophilic strains (*A. tundrae* and *A. paludosum*) encoded for twice as many cold shock proteins (4 total) than the other *Acetobacterium* strains (1-2 total).

Other minor variations were observed in the psychrophilic strains, in particular the sodium-dependent ATPase and 1,2-propanediol operon. The subunits of the sodium-dependent ATPase involved in pore formation and proton transport were less well conserved than the catalytic subunits, perhaps indicative of the varying membranes across *Acetobacterium* (Supplemental Data). Minor variability in the 1,2-propanediol operon structures of three psychrophilic strains (*A. fimetarium, A. paludosum,* and *A. tundrae*) was evident, as a heme-binding protein (pfam03928) located between *pduO* and *pduP* was missing.

One adaptation to survival in cold environments psychrophiles employ is to increase protein flexibility and stability [98]. As a result, psychrophiles often contain a higher proportion of hydrophobic amino acid residues, such as alanine and glycine [99, 100]. The psychrophilic strains are the only sequenced *Acetobacterium* strains to lack an alanine degradation pathway and glycine cleavage system. One possible explanation for the loss of these pathways in the psychrophilic *Acetobacterium* strains is an increased utilization of alanine and glycine in cold-adapted protein synthesis [99]. Interestingly, the alanine degradation pathway encodes for alanine dehydrogenase, which may also play an important role in NH_4_ assimilation [101]. Furthermore, glutamate dehydrogenase, another enzyme shown to be important for nitrogen assimilation, was also absent from the psychrophilic strains (Supplemental File). Thus, the psychrophilic strains may have adapted to obtain nitrogen from alternative sources, using an alternative pathway.

To further determine what functional genome attributes may separate the psychrophilic *Acetobacterium* species from their non-psychrophilic counterparts, we identified gene families unique to the psychrophilic species (Supplemental File). Our analyses revealed 40 annotated gene families found only in the psychrophilic clade, and included a calcium-translocating P-type ATPase, sodium:proton antiporter, cupin, ribulose 1,5-bisphosphate carboxylase, and SDR family oxidoreductase, among others (Supplemental File). The presence of a unique calcium-translocating P-type ATPase and sodium:proton antiporter suggest modified abilities to transport calcium and sodium. The 1,5-bisphosphate carboxylase (RuBisCO) and cupin may be involved in the methionine salvage pathway, which has been observed in other bacteria [102, 103]. SDR family oxidoreductases have a wide range of activities, including metabolism of amino acids, cofactors, carbohydrates, and may play a role in redox sensing [104]. More work is needed to determine the exact function each protein plays in the psychrophilic strains, but these findings can be used as a framework for targeted functional analyses in future studies to determine specific roles these unique proteins play in the metabolisms of psychrophilic *Acetobacterium* strains and their persistence in cold environments.

## Conclusions

Acetogens are a phylogenetically diverse group of microorganisms capable of converting CO_2_ into acetate. *Acetobacterium woodii* has been used as a model organism to study the WLP and accessory components required for energy conservation in acetogens lacking cytochromes ([23], and references therein). In this study we sequenced four *Acetobacterium* isolates from the culture collection (ATCC) (*A. fimetarium, A. malicum, A. paludosum,* and *A. tundrae*). Using comparative genomics, we examined the functional potential of the available 13 *Acetobacterium* genomes to shed light on the diverse genome attributes, gene arrangement and architecture, and potential metabolic capabilities of the *Acetobacterium* genus. Using the type strain of the genus (*A. woodii*) as a framework for pathway identification and comparison, we found the common and conserved pathways included the WLP and accessory components [electron bifurcating hydrogenase, ATP synthase, Rnf complex, ferredoxin, and electron transfer flavoprotein (ETF)], glycolysis/gluconeogenesis, and 1,2-propanediol (Figure 5). Divergent metabolisms found in a subset of genomes included caffeate reduction, 2,3-butanediol oxidation, ethanol oxidation, alanine metabolism, glycine cleavage system, and methanol oxidation. Notably, the psychrophilic strains encode for unique amino acid transport and utilization, and ion transport, which may have evolved for survival in low-temperature environments, in addition to post-transcriptional modifications [32]. Members of the *A. wieringae* clade encode for unique diguanylate cyclases and a unique methylene-THF reductase, which may aid in attachment and colonization to solid surfaces. Overall, the comparative genomic analysis performed on the *Acetobacterium* genus provides a framework to understand the conserved metabolic processes across the genus, as well as identifying divergent metabolisms that can be exploited for targeted biotechnological applications using *Acetobacterium* strains.

**Figure 5.**
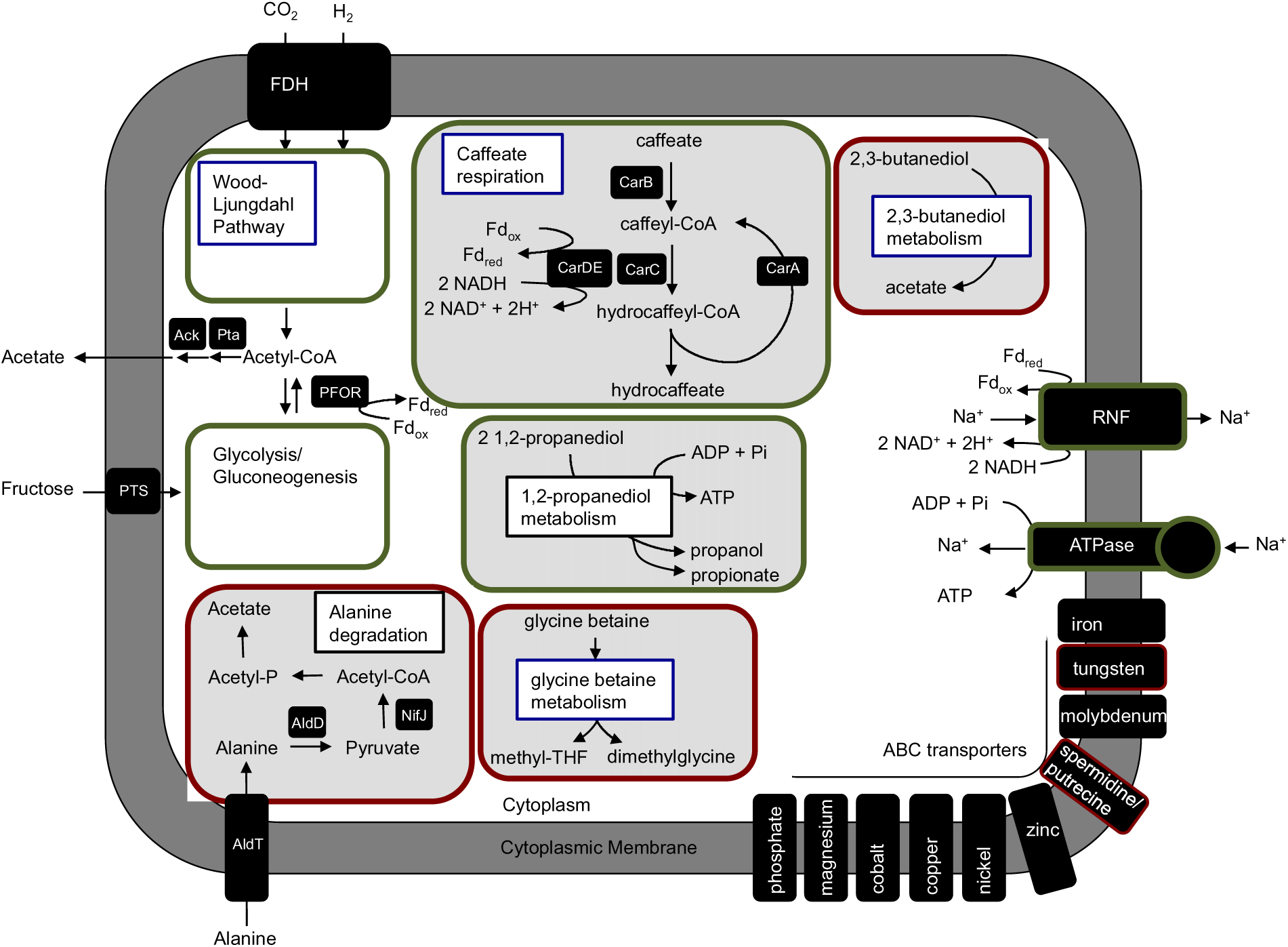
Metabolic overview of the *Acetobacterium* genus. Major metabolic pathways have been biochemically and genetically verified in *Acetobacterium woodii*, and predicted metabolisms are based upon the presence of each operon and similarity of predicted amino acid sequences. Pathways and accessory components framed in green are encoded by all sequenced *Acetobacterium* strains. Pathways denoted in red are found in only a subset of *Acetobacterium* strains. Metabolic pathways denoted in blue (*e.g.* glycine betaine, caffeate, and 2,3-butanediol) are linked to the Wood-Ljungdahl pathway.

## Authors and Contributors

DER performed analyses, analyzed data, and wrote the manuscript. CWM analyzed data and wrote the manuscript. HM, RSN, and DG edited the manuscript.

## Conflict of interest

The authors declare no conflict of interest.

## Funding

Funding was provided by the U.S. Department of Energy, Advanced Research Project Agency–Energy (award DE-AR0000089).

## Acknowledgement

This work was performed in support of the US Department of Energy’s Fossil Energy CO_2_ Utilization Research Program. The Research was executed through the NETL Research and Innovation Center’s CO_2_ Utilization Technologies FWP. Research performed by Leidos Research Support Team staff was conducted under the RSS contract 89243318CFE000003.

## Disclaimer

This work was funded by the Department of Energy, National Energy Technology Laboratory, an agency of the United States Government, through a support contract with Leidos Research Support Team (LRST). Neither the United States Government nor any agency thereof, nor any of their employees, nor LRST, nor any of their employees, makes any warranty, expressed or implied, or assumes any legal liability or responsibility for the accuracy, completeness, or usefulness of any information, apparatus, product, or process disclosed, or represents that its use would not infringe privately owned rights. Reference herein to any specific commercial product, process, or service by trade name, trademark, manufacturer, or otherwise, does not necessarily constitute or imply its endorsement, recommendation, or favoring by the United States Government or any agency thereof. The views and opinions of authors expressed herein do not necessarily state or reflect those of the United States Government or any agency thereof.

## Notes

https://narrative.kbase.us/narrative/ws.53630.obj.1

