## Supplemental Text, Tables, and Figures for "The defining genomic and predicted metabolic features of the Acetobacterium genus"

Daniel E. Ross

**Table of contents:**

1. **Supplementary Results and Discussion**
   a. **Insight into the placement of the most incomplete *Acetobacterium* MAGs.**
   b. **Effect of genome completeness on core genome partitioning.**
   c. **KEGG modules identified in *Acetobacterium* core genome.**
   d. **Pan-genome analysis of the *Eubacteriaceae* family.**
   e. **Variability in Cluster I of the Wood-Ljungdahl pathway encoding the formate dehydrogenase.**
   f. **Variability in the number of ferredoxins encoded by *Acetobacterium* genomes.**
   g. **Variability in electron transfer flavoprotein (ETF)**
   h. **Potential alternative mechanism for ethanol oxidation.**
   i. **Membrane transport.**
   j. **Amino acid transport and biosynthesis**
   k. **Glycine betaine metabolism.**
   l. **Unique methylene-THF reductase of the *A. wieringae* clade**.
   m. **Phylogenetic analysis and genome architecture surrounding the unique diguanylate cyclases (DGC) of the *A. wieringae* clade.**
   n. **Motility.**

**Supplemental Table 1. Genome statitstics of four sequenced *Acetobaterium* isolates.**

**Supplemental Figure 1. Variability in genome architecture of WLP Cluster I.**

**Supplemental Figure 2. Electron bifurcating hydrogenase.**

**Supplemental Figure 3. Evidence for Na^+^ binding motif in *c*-subunits of *Acetobacterium* F_1_F_0_- ATP synthase complex.**

**Supplemental Figure 4. *Acetobacterium* pan-genome partitioning at various gene family clustering cutoff values.**

**Supplemental Figure 5. Pan-phylogeny of 29 Eubacteriaceae genomes.**

**Supplemental Figure 6. Core phylogeny of 29 Eubacteriaceae genomes.**

**Supplemental Figure 7. Phylogenetic analysis of 5 uniqe diguanylate cyclases from the *A. wieringae* clade.**

**RESULTS**

**Insight into the placement of the most incomplete *Acetobacterium* MAGs within the *Acetobacterium* genus.** Ideally whole genome comparative genomics is performed on complete genomes, enabling the most accurate assessment of which genes (and predicted protein sequences) are conserved across a group of genomes [1, 2]. In order to obtain the most complete and accurate pan-genome statistics, only genomes >98% were utilized for pan-genome analysis (Figure 2). Here we show the phylogenomic placement of the three most incomplete MAGs; *A.* sp. UBA6819 (94% complete, missing 11 marker genes), *A.* sp. UBA5558 (85% complete, missing 31 marker genes), and *A.* sp. UBA11218 (57% complete, missing 93 marker genes). Pan-genome analyses of all sequenced stains irrespective of completeness level, including *A.* sp. UBA6819, *A.* sp. UBA5558, and *A.* sp. UBA11218, revealed pan-genome and core genome clustering of *A.* sp. UBA5558, and *A.* sp. UBA11218 within the A. *wieringae* clade (Supplemental Figure 5 and 6), and *A.* sp. UBA6819 with *A.* sp. KB-1. Phylogenomic placement of the other strains did not change when including all *Acetobacterium* strains. The only observable difference was the partitioning of the core, accessory, and unique gene families, with a decrease in the core genome. These findings support ANI and AAI values, where *A.* sp. UBA6819 was most closely related to *A.* sp. KB-1, and *A.* sp. UBA5558, and *A.* sp. UBA11218 were >95% similar to *A. wieringae* and *A.* sp. MES1.

**Effect of gene family clustering cutoffs on core genome partitioning.** Another variable in our pan-genome analyses was the cutoff value used for clustering similar sequences into gene families. The default cutoff for the pan-genome software utilized here (BPGA) is 50% [3]. To determine how the clustering cutoff affects pan-genome partitioning (*i.e.* clustering of gene families into core, accessory, and unique categories), we ran our analyses on a defined set of genomes while varying the clustering cutoff for each run. We examined how the number of core genome families changes with varying clustering cutoffs. Utilizing a range of cutoff values from 5% to 99% the highest number of core gene families were found with clustering cutoffs between 30 to 50% (Supplementary Figure 4).

**KEGG modules identified in *Acetobacterium* core genome.** Examination of the core genome (11 strains) using a 50% clustering cutoff revealed complete KEGG pathway modules for phosphate acetyltransferase-acetate kinase pathway (acetyl-CoA to acetate), nitrogen fixation (nitrogen to ammonia), pyruvate oxidation (pyruvate to acetyl CoA), citrate cycle (oxaloacetate to 2-oxoglutarate), glycogen biosynthesis (glucose-1P to glycogen), nucleotide sugar biosynthesis (glucose to UDP-glucose), dTDP-L rhamnose biosynthesis, fatty-acid biosynthesis (initiation and elongation), inosine monophosphate biosynthesis (PRPP + glutamine to IMP), F-type ATPase, adenine ribonucleotide biosynthesis (IMP to ADP,ATP), prokaryotes, DNA polymerase III complex, D-methionine transport system, cobalt/nickel transport system, putative ABC transport system, cell division transport system, ABC-2 type transport system, PTS system (fructose-specific II component), and Sec (secretion) system (Supplemental File).

**Pan-genome analysis of the *Eubacteriaceae* family.** Acetogens are phylogenetically diverse and have been found in over 20 different phyla to date [4]. The *Acetobacterium* genus contains only acetogens, but the genus is part of the *Eubacteriaceae* family that contains both acetogens and non-acetogens. We performed pan-genome analyses to determine the predicted protein sequences that are common to the *Eubacteriaceae* family. We determined the completeness of each genome using the *Eubacteriaceae* marker gene set from CheckM [5]. Of the available genomes, only 29 were above the desired completeness cutoff of 95%. We examined the pan-genome of the 29 most complete genomes from the *Eubacteriaceae* family to determine the common pathways of the family and phylogenomic placement of *Acetobacterium* within the family (Supplementary Figure 5 and 6). While the number of core gene families decreased significantly (from 1406 to 44), the trend was the same—the highest number of core gene families were observed when using cutoffs between 30 and 50% (Supplemental Figure 4). The *Eubacteriaceae* core genome (50% gene family cutoff) consisted of 44 gene families, the majority of which were mostly related to genetic information and processing—specifically, translation (ribosome, aminoacyl-tRNA biosynthesis); folding, sorting, and degradation (RNA degradation, protein transport); and replication and repair (nucleotide excision repair and homologous recombination) (Supplemental File).

**Variability in Cluster I of the Wood-Ljungdahl pathway encoding the formate dehydrogenase.** Formate dehydrogenase (FdhF1) is the first protein encoded by Cluster I in *Acetobacterium woodii*. The gene content and arrangement in Cluster I is varied amongst the *Acetobacterium* genus and only the psychrophilic strains encode for the *A. woodii*-type hydrogen dependent CO_2_ reductase (HDCR) (Supplemental Figure 1). Examination of the remaining *Acetobacterium* genomes for this protein revealed a best hit to a protein annotated as an NAD-dependent formate dehydrogenase alpha subunit, which was quite divergent at the amino acid level, with a percent identity and coverage between *A. woodii* FhdF1 and *A.* sp. MES1 NAD-dependent formate dehydrogenase of 39% and 58%, respectively.

The genome architecture surrounding the NAD-dependent formate dehydrogenase had identical synteny for various groups of strains. For example, the *A. wieringae* clade and *A. dehalogenans* Cluster I included NAD-reducing hydrogenase subunit HoxE (EC 1.12.1.2), electron-bifurcating [FeFe]-hydrogenase subunit C (2Fe2S protein), electron-bifurcating [FeFe]-hydrogenase subunit B (NAD+ reductase, ferredoxin reductase), NAD-dependent formate dehydrogenase alpha subunit, formylmethanofuran dehydrogenase associated protein FmdE/molybdopterin molybdenum transferase (EC 2.10.1.1), and formate dehydrogenase chain D (EC 1.2.1.2) (Supplemental File). The NAD-dependent formate dehydrogenase encoded by *A.* sp. KB-1 and *A.* sp. UBA6819 had similar genome architecture immediately upstream with a cadmium efflux protein, and heavy-metal transporting ATPase nearby (Supplemental File).

**Variability in the number of ferredoxins encoded by *Acetobacterium* genomes.**

Ferredoxins are soluble cellular redox compounds and in acetogens, play a major role in the generation of a transmembrane ion gradient for ATP generation during CO_2_ fixation [6]. Of the electron carriers involved in the WLP, only ferredoxins have been demonstrated to provide the reducing potential required for the reduction of CO_2_ to CO in the carbonyl branch. The genome of *Acetobacterium woodii* encodes for 17 ferredoxins, the most of any *Acetobacterium* genome to date. The number of ferredoxins encoded by the other *Acetobacterium* genomes ranged from 6 (*A.* sp. KB-1 and *A.* sp. UBA6819) to 16 (*A. bakii*). Specifically, *A. malicum* and *A. dehalogenans* encode for 12 ferredoxins, members of the *A. wieringae* clade encoded for 8-10, and *A.* sp. KB-1 and *A.* sp. UBA6189 encoded for six. The psychrophilic clade was more varied, as *A. fimetarium* encoded for 8, while *A. paludosum* encoded for 11, *A. tundrae* encoded for 13, and *A. bakii* encoded for 16. The overall variability in the number of ferredoxins encoded by each genome may be a casualty of loss of other things in the genome or an indication that it has less or more specific electron transport functions to carry out.

**Variability in electron transfer flavoprotein (ETF) copy number.** Electron transfer flavoprotein (Etf) is an electron carrier in the oxidation of NADH in Rnf complex-mediated sodium transport in *Acetobacterium woodii* [7]. *A. woodii* encodes for two separate EtfAB modules. One is adjacent to CarABC and part of the caffeate reduction operon (CarDE), while the other is directly upstream of glycolate dehydrogenase, L-lactate permease, and D-mannotate dehydrogenase (Supplementary File) and involved in lactate metabolism [8]. Examination of the remaining 12 genomes revealed that, with the exception of *A.* sp. UBA5558, all genomes encode for at least one copy of EtfAB. Specifically, *A. tundrae* encodes for four copies of EtfAB; *A. bakii* and *A. paludosum* encode for three copies; *A. woodii*, and *A. dehalogenans* each encode for two copies; and the remaining strains have a single copy (Supplementary File). The gene architecture downstream of the single EtfAB module encoded by all strains (except *A.* sp. UBA5558) was identical, and included machinery for lactate metabolism (Supplementary File). The four EtfAB modules of *A. tundrae* weren’t well conserved, with amino acid identities EtfA ranging from 39% to 55% and EtfB ranging from 43-48%.

**Potential alternative mechanism for ethanol oxidation.** *Acetobacterium woodii* utilizes a bifunctional acetaldehyde-CoA/alcohol dehydrogenase (AdhE) and encodes for 7 alcohol dehydrogenases for ethanol oxidation [9]. The only other *Acetobacterium* strains that encode for AdhE were *A.* sp. KB-1 and *A.* sp. UBA6819 (87% identity, 95% coverage)*.* Examination of the *A.* sp. MES1 MAG revealed an alcohol dehydrogenase upstream of a tungsten-containing aldehyde:ferredoxin oxidoreductase. Other *Acetobacterium* genomes harbored a similar alcohol dehydrogenase, with the exception of *A. woodii, A. paludosum,* and *A. tundrae*. Genome architecture surrounding the alcohol dehydrogenase and tungsten-containing aldehyde:ferredoxin oxidoreductase was similar for many MAGs and included an alcohol dehydrogenase, hypothetical protein, 2-oxoglutarate oxidoreductase alpha and beta subunits, hypothetical protein or superoxide reductase, alcohol dehydrogenase, hypothetical protein, tungsten-containing aldehyde:ferredoxin oxidoreductase, and a hypothetical (Supplementary File). *A.* sp. MES1, *A.* sp. UBA5558, and *A.* sp. UBA5834 contained a CRISPR array downstream of the last hypothetical protein. All other genomes, with the exception of *A. paludosum* and *A. tundrae,* contained a variety genes surrounding the alcohol dehydrogenase, including an acetyltransferase; fumarate reductase flavoprotein; PTS system, glucose-specific IIC component / PTS system, glucose-specific IIB component (EC 2.7.1.69); and two chromate transport proteins (Supplementary File).

**Membrane transport.** Each *Acetobacterium* genome encodes for numerous ABC transporters, specific to heavy metals (*e.g.,* iron, tungsten, molybdenum, zinc, nickel), oligopeptides, branched chain amino acids, dipeptides, phosphate and cations (*e.g.,* copper, nickel, cobalt, and magnesium). *A.* sp. UBA6819 encoded for the most ABC transporters (190), while *A.* sp. UBA5558 encoded for the least (83). Interestingly, three psychrophilic strains (*A. fimetarium, A. paludosum,* and *A. tundrae*) were missing ABC transporters specific for spermidine/putrescine, and hydroxymethyl pyrimidine (Supplemental File). Spermidine and putrescine are polyamines that bind to and modulate the function of intracellular nucleic acids and ATP [10], play a role in biofilm formation [11], and combat oxidative and nitrosative stress. It is unknown why these psychrophilic strains lack this transporter, but spermidine and putrecine are involved in the methionine salvage pathway. Despite the lack of spermidine/putrecine-specific ABC transporters, the psychrophilic strains encode for spermidine biosynthetic enzymes S-adenosylmethionine decarboxylase (EC 4.1.1.50), arginine decarboxylase (EC 4.1.1.19), spermidine synthase (EC 2.5.1.16), carboxynorspermidine synthase (EC 1.5.1.43), and carboxynorspermidine decarboxylase (EC 4.1.1.96).

**Amino acid transport and biosynthesis.** Pathways specific to amino acid biosynthesis were conserved across the *Acetobacterium* genus, including cysteine biosynthesis (serine to cysteine), proline biosynthesis (glutamate to proline), and histidine biosynthesis (PRPP to histidine). Transport systems for polar amino acids (serine, threonine, cysteine, proline, asparagine, and glutamine), and branched-chain amino acids (leucine, isoleucine, and valine) were also present and well conserved.

**Glycine betaine metabolism.** Glycine betaine (GB) can be metabolized under anoxic conditions by acetogenic bacteria and examination of this process in *Acetobacterium woodii* revealed a gene cluster responsible for GB utilization as a carbon and energy source [12]. In *Acetobacterium bakii*, the GB operon is more highly expressed at cold temperatures during heterotrophic growth on fructose [13] and thus may play an important role in growth at low temperatures. The gene cluster is comprised of 5 genes, and includes *mttA2, opuD2, mttB10, mttC6,* and *opuD3* (Awo_c07520-Awo_c07560). The *mttA2* gene encodes for a methyltransferase, *opuD2 and opuD3* encode for choline/carnitine/betaine transporters, *mttB10* encodes for a methyltransferase, and *mttC6* encodes for a corrinoid protein. All sequenced *Acetobacterium* strains, with the exception of *A.* sp. UBA5558 contain this operon and each protein encoded by this operon has a high sequence similarity to the GB operon in *A. woodii* (Supplementary File). Interestingly, the choline/carnitine/betaine transporter OpuD3 is divergent (42% identity, 62% positives) in three psychrophilic strains (*A. fimetarium, A. paludosum,* and *A. tundrae*), suggesting modified transport of GB or a system tuned to environmental stressors that allows for optimal use of GB, either as a substrate or as an osmoprotectant [14].

**Unique methylene-THF reductase of the *A. wieringae* clade**. One of the amino acid sequences unique to the *A. wieringae* clade was annotated as a methylenetetrahydrofolate reductase (Methylene-THF reductase). Examination of the MAGs containing this predicted protein revealed the methylene-THF reductase was located downstream of a sodium/sulfate symporter in all strains. In *A.* sp. MES1, *A.* sp. UBA5558, and *A. sp.* UBA5834, it was located upstream of an Fis family transcriptional regulator and a CO dehydrogenase/CO-methylating acetyl-CoA synthase complex (subunit beta). The methylene-THF reductase was found at the end of a contig in *A. wieringae*. Interestingly, the Fis family transcriptional regulator and a CO dehydrogenase/CO-methylating acetyl-CoA synthase complex (subunit beta) was found at the end of a different contig. The end of each contig contained a repeat region that overlapped by 23 bp. Based upon these findings, it is likey that the genome architechture surroundning the methylene-THF reductase in *A. wieringae* is identical to that of *A.* sp. MES1, *A.* sp. UBA5558, and *A. sp.* UBA5834.

**Phylogenetic analysis and genome architecture surrounding the unique diguanylate cyclases (DGC) of the *A. wieringae* clade.** Diguanylate cyclases (DGCs) control intracellular levels of cyclic-di-GMP in concert with EAL- or HD-GYP-domain containing phosphodiesterases (PDE, hydrolases) that have opposing activities [15]. *A. wieringae, A.* sp. MES1, and *A. dehalogenans* encode for 45, 51, and 51 DGCs, respectively, and no other strain encodes for more than 35. Pan-genome analysis revealed five DGCs unique to the *A. wieringae* clade. The five DGCs were examined individually and the best BLAST hits to the other MAGs were used to generate a tree based upon aligned amino acid sequences. Tree generation revealed succinct clustering of the *A. wierinage* clade DGCs apart from the remaining *Acetobacterium* DGCs (Supplementary Figure 7).

The first unique DGC (representative accession: OFV69692.1) was annotated as a cyclic di-GMP phosphodiesterase response regulator RpfG and was surrounded by three methyl-accepting chemotaxis proteins. The second (WP_070371128) was annotated as a diguanylate cyclase and is upstream of a branched chain amino acid transport system, and in all MAGs but *A. wieringae*, the DGC was downstream of a CRISPR-Cas array. The third (WP_070372088.1) was annotated as a DGC and located directly upstream of a methyl-accepting chemotaxis protein. The fourth (WP_070370773) was annotated as a DGC and located directly downstream of two lipid A export proteins (MsbA), and directly upstream of a transcriptional regulator (YpdB) and a signal transduction histidine kinase (CheA). The fifth (WP_070371683.1) was annotated as a DGC and located directly upstream of enzymes involved in oxidative stress (H_2_O_2_ scavenging and resistance to reactive oxygen species (ROS)) and include an alkyl hydroperoxide reductase subunit-C, dehydrosqualene desaturase, phytoene synthase, phytol kinase, and a zinc-containing alcohol dehydrogenase.

**Motility.** Motility in *Acetobacterium* arises from the activity of one or two sub-terminal flagella [16]. The thirteen *Acetobacterium* genomes contain 27-34 features related to various flagella function and include flagellar biosynthesis proteins FlhAB and FliPQRS, flagellar basal-body rod proteins FlgBCF, flagellar hook-associated proteins FlgKL, flagellar hook protein FlgE, flagellar hook-length control protein FliK, flagellar hook-basal body complex protein FliE, flagellar M-ring protein FliF, flagellar motor rotation proteins MotAB, flagellar motor switch proteins FliGMN, flagellin protein FlaA, RNA polymerase sigma factor for flagellar operon, and a flagellum-specific ATP synthase FliI.

**Supplemental Tables and Figures.**

**Supplemental Table 1**. Genome statitstics of four sequenced *Acetobaterium* isolates.

| **Genome** | ***A. fimetarium*** | ***A. malicum*** | ***A. paludosum*** | ***A. tundrae*** |
| --- | --- | --- | --- | --- |
| **Accession number** | WJBC00000000 | WJBE00000000 | WJBD00000000 | WJBB00000000 |
| **Total bp** | 3,240,781 | 4,079,435 | 3,690,621 | 3,560,849 |
| **% reads mapped to assembly** | 99.93 | 99.80 | 99.37 | 99.46 |
| **Predicted genes** | 3,034 | 4,071 | 3,427 | 3,383 |
| **G + C %** | 44.69 | 43.73 | 40.05 | 39.66 |
| **Contigs > 500 bp** | 81 | 72 | 66 | 70 |
| **Contigs > 1000 bp** | 74 | 60 | 54 | 63 |
| **Contig N50** | 117,876 | 224,498 | 160,493 | 160,928 |
| **Largest contig** | 304,900 | 532,947 | 402,626 | 330,645 |
| **Genome completion** | 98.57% | 99.29% | 99.29% | 99.29% |
| **Contamination** | 0.71% | 1.43% | 2.14% | 1.43% |
| **5S, 16S, and 23S** | yes | yes | yes | yes |
| **tRNAs (amino acids encoded)** | 56 (20) | 54 (20) | 58 (20) | 58 (20) |

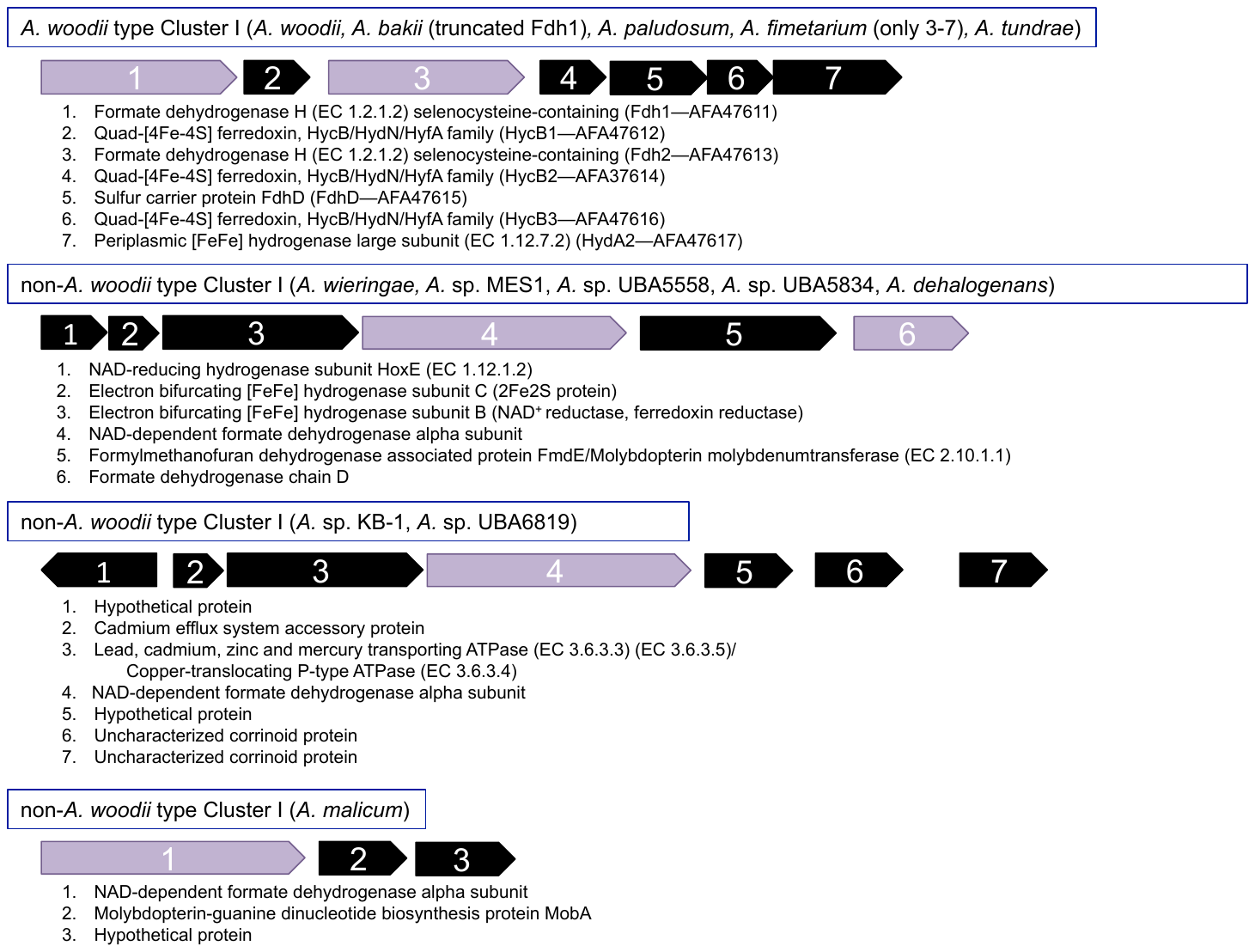

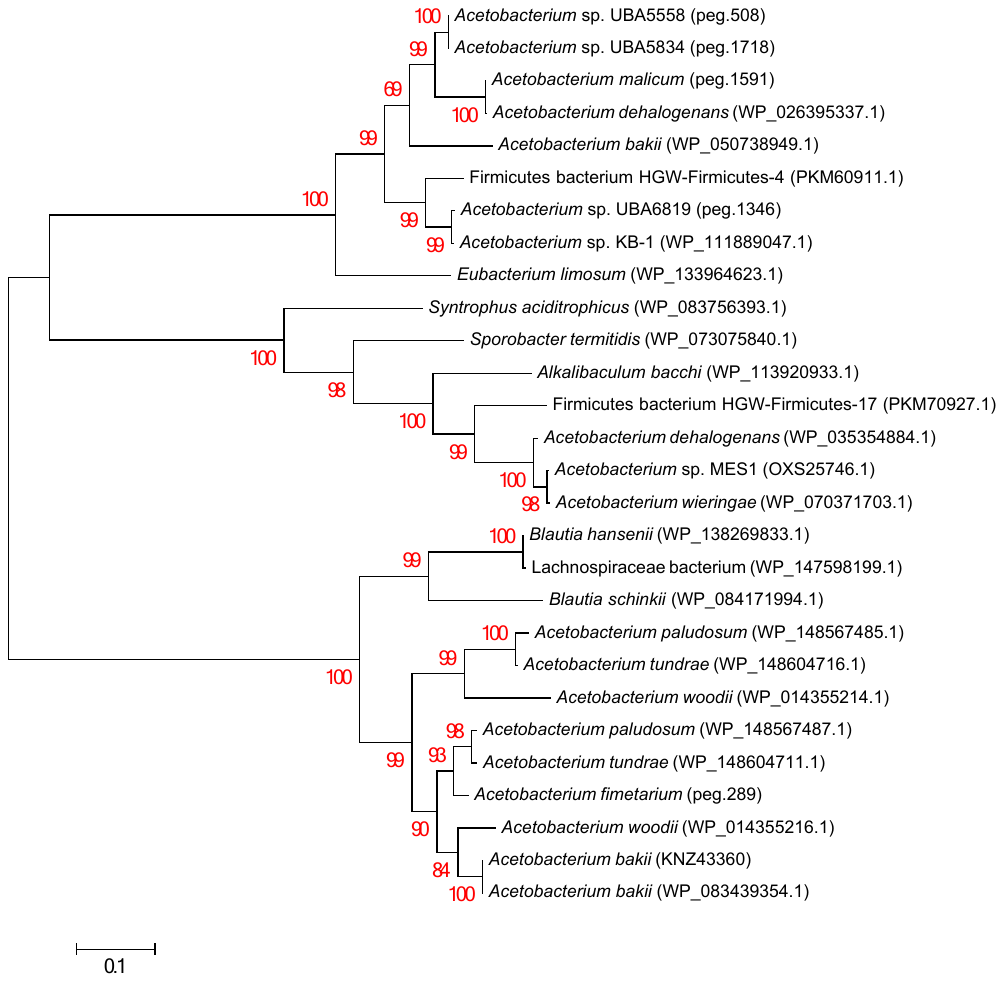

**Supplemental Figure 1. Variability in genome architecture of WLP Cluster I.** A) Gene arrangement of Cluster I of the Wood-Ljungdahl Pathway. The formate dehydrogenase subunits are denoted in purple. B) Phylogenetic tree of FdhF1 and NAD-dependent formate dehydrogenases from *Acetobacterium* and closely-related proteins.

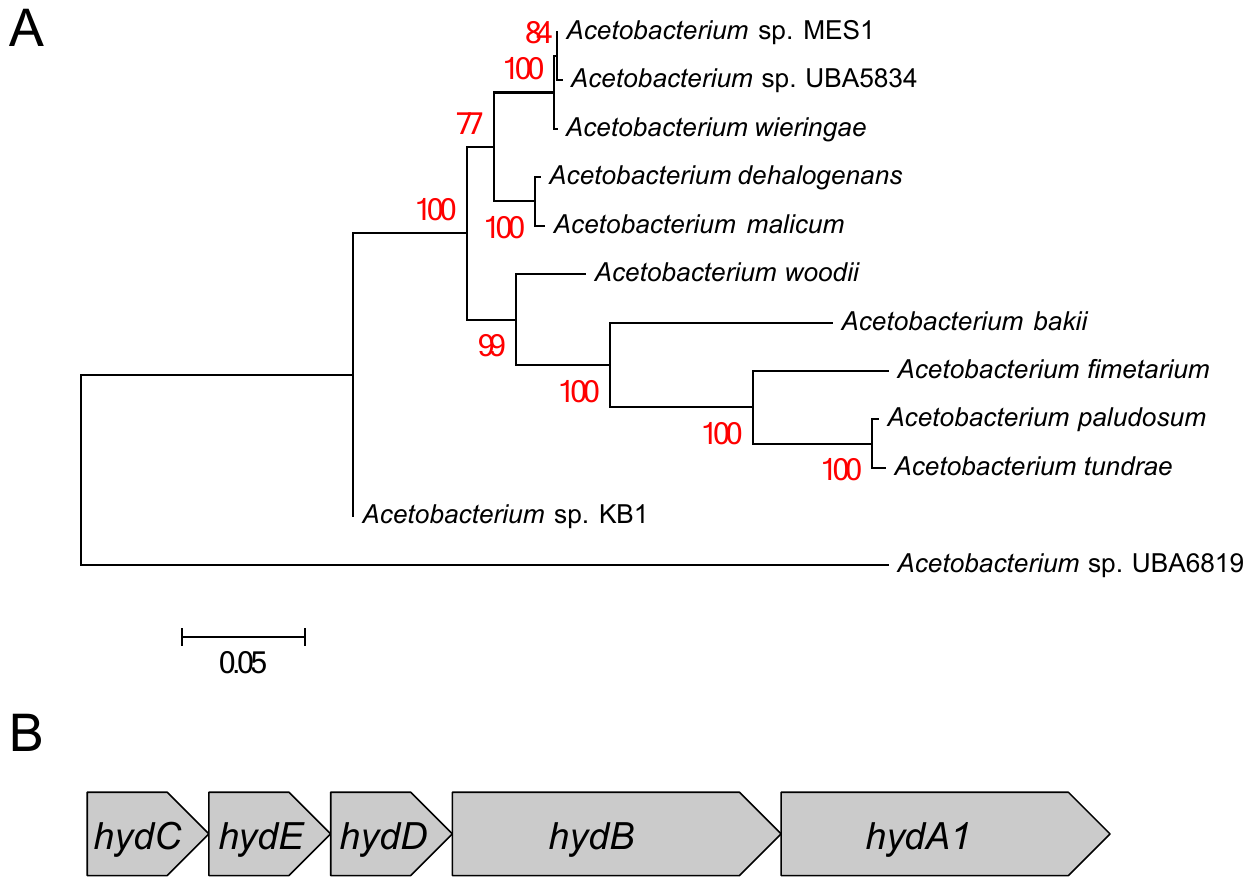

**Supplemental Figure 2.** **Electron bifurcating hydrogenase**. A) Phylogenetic tree of the HydABDEC predicted protein sequences. Protein sequences of HydA, HydB, HydD, HydE, and HydC were concatenated and clustered with MUSCLE [17] and the phylogenomic tree was constructed with the Maximum Likelihood method using the Bootstrap method with 1000 replications for test of phylogeny. The substitution method was Jones-Taylor-Thornton with uniform rates among sites. Clustering and tree construction were performed with MEGA 6.06 [18]. MAG *A.* sp. UBA5558 was not included in the analysis due to the low sequence similarity of HydA1 (41% identity, 58% coverage) and HydE (24% identity, 46% coverage), most likely due to the incompleteness of the MAG. B) Gene arrangement of the hydrogenase operon.

**Supplemental Figure 3. Evidence for Na^+^ binding motif in *c*-subunits of *Acetobacterium* F_1_F_0_-type ATP synthase complex.** Three *c-*subunits from *Acetobacterium woodii* (Awo_c02160 – c02180) were used to query the remaining 12 *Acetobacterium* genomes in RAST. All strains encode for at least two subunits with binding motifs specific for Na^+^ (denoted in red). Three of the psychrophilic strains contain multiple amino acid residues at varying positions surrounding the Na^+^ binding motif that differ from all other sequenced *Acetobacterium* strains. The psychrophilic strains also only encode for two *c*-subunits, while the others encode for three.

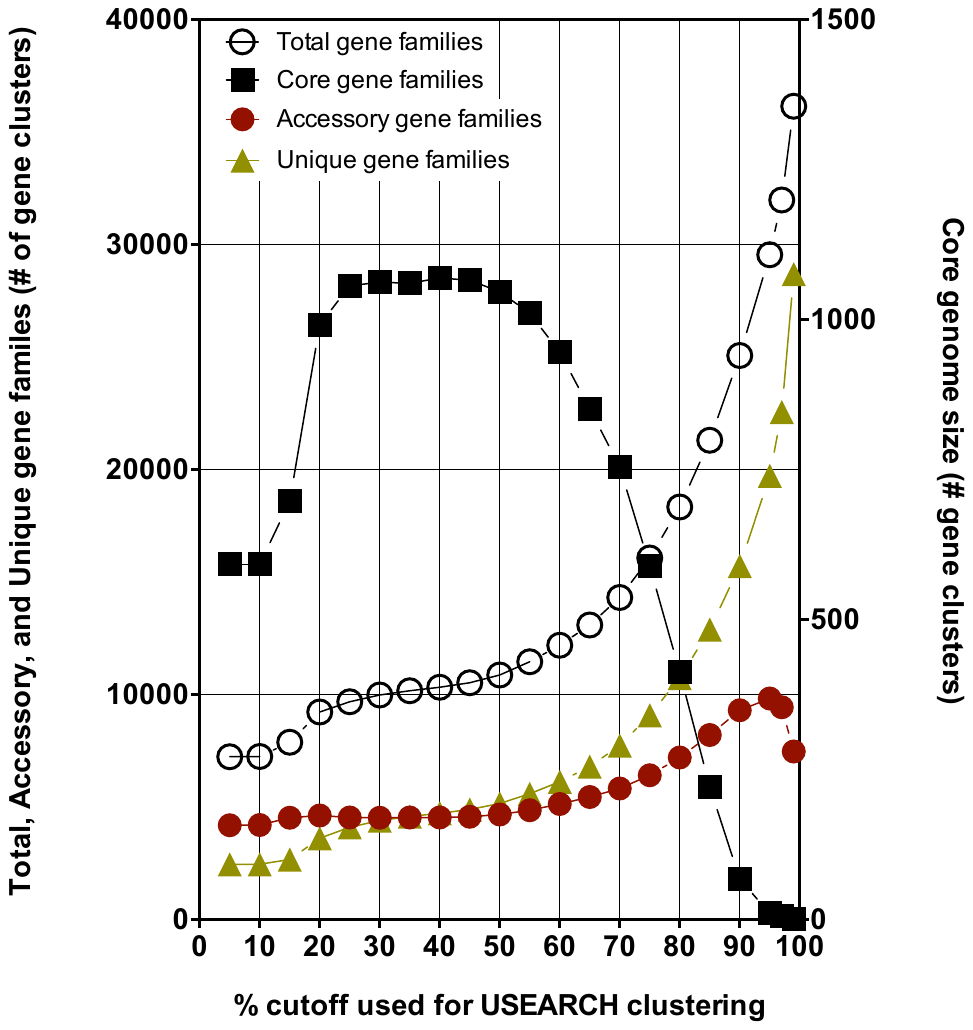

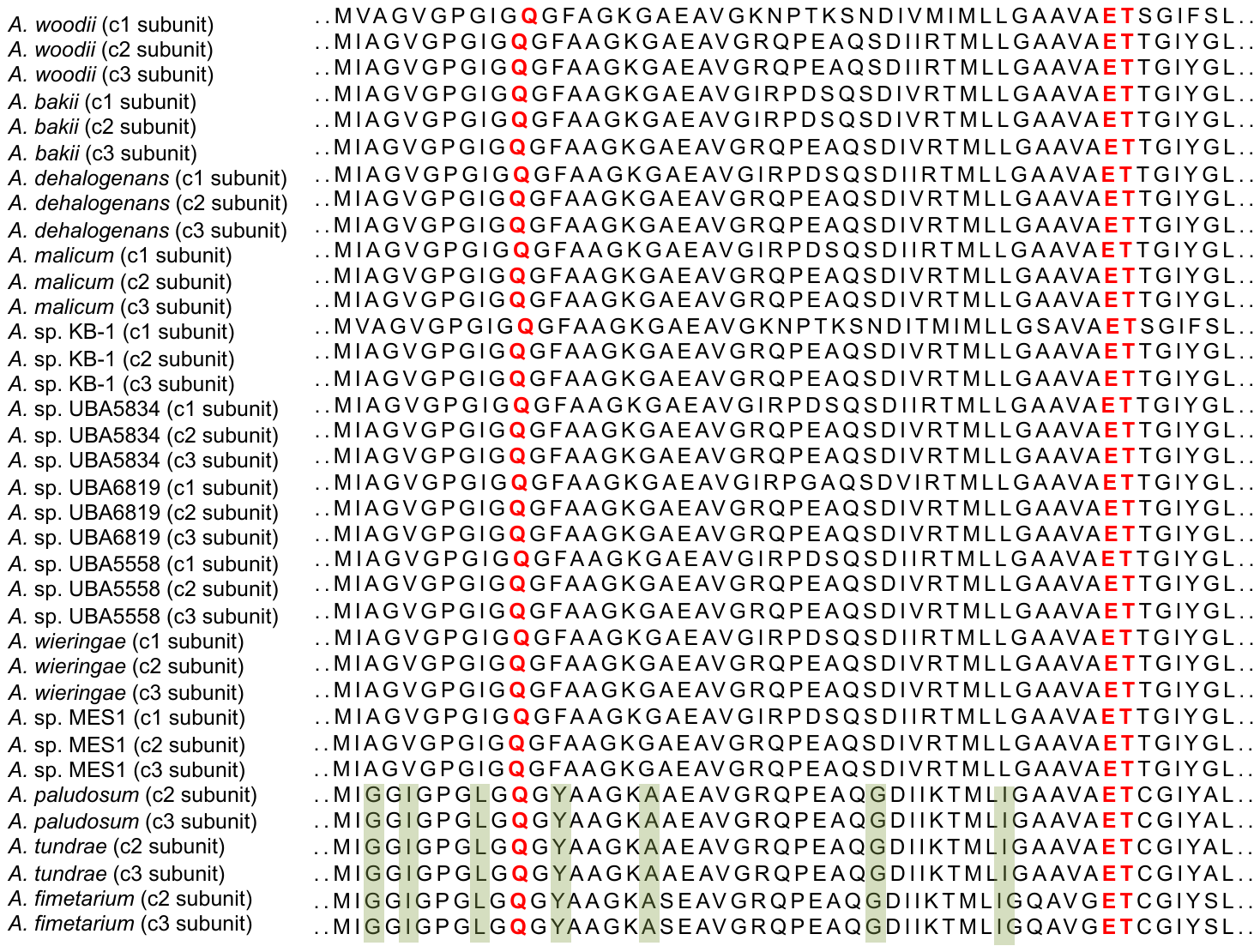

**Supplemental Figure 4. *Acetobacterium* pan-genome partitioning at various gene family clustering cutoff values.** The effect of varying clustering cutoff values on the pan-genome partitioning. The total number of core gene families plateaued between 25 and 45%, and decreased linearly from 60-90%.

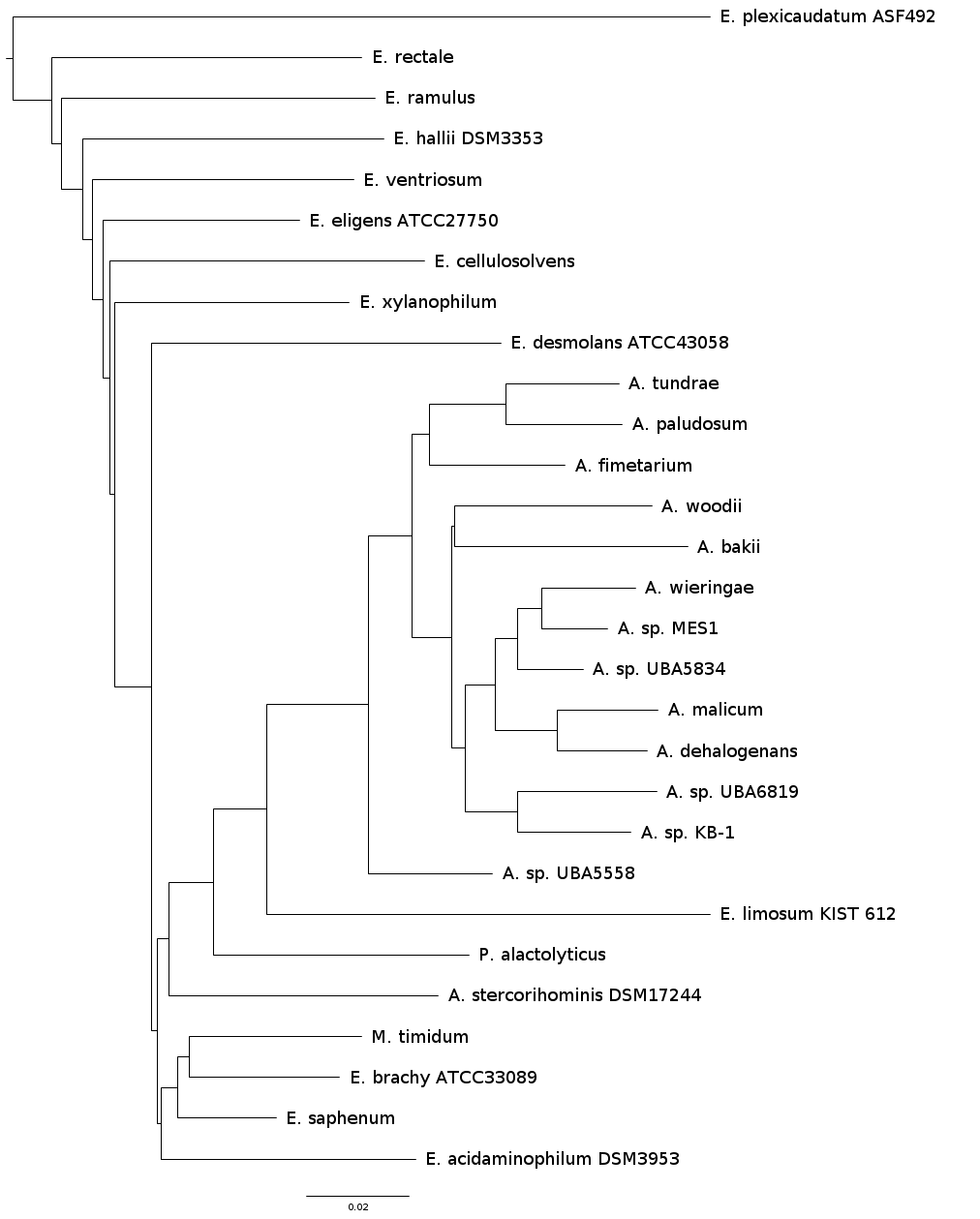

**Supplemental Figure 5. Pan-phylogeny of 29 Eubacteriaceae genomes.**

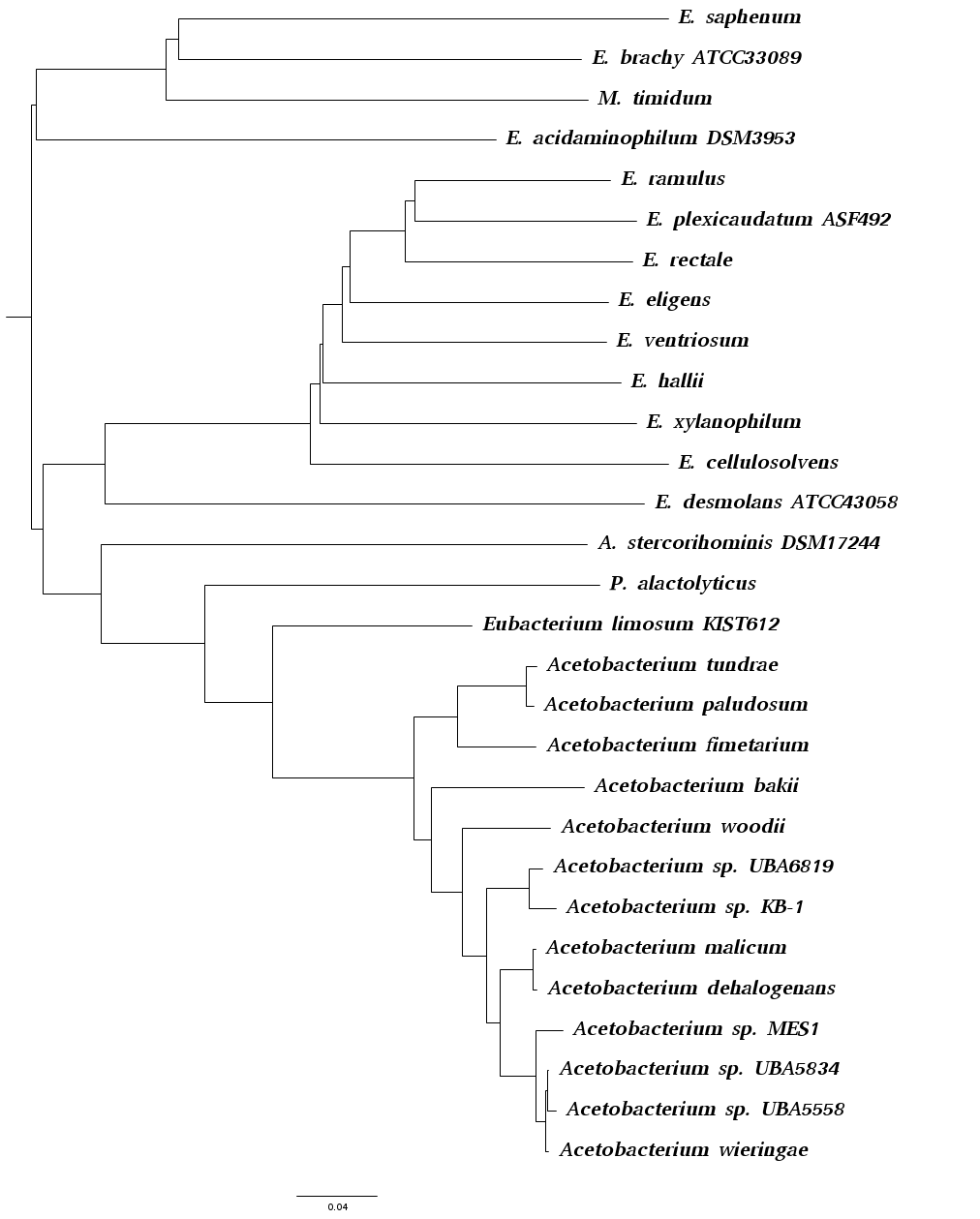

**Supplemental Figure 6. Core phylogeny of 29 Eubacteriaceae genomes.**

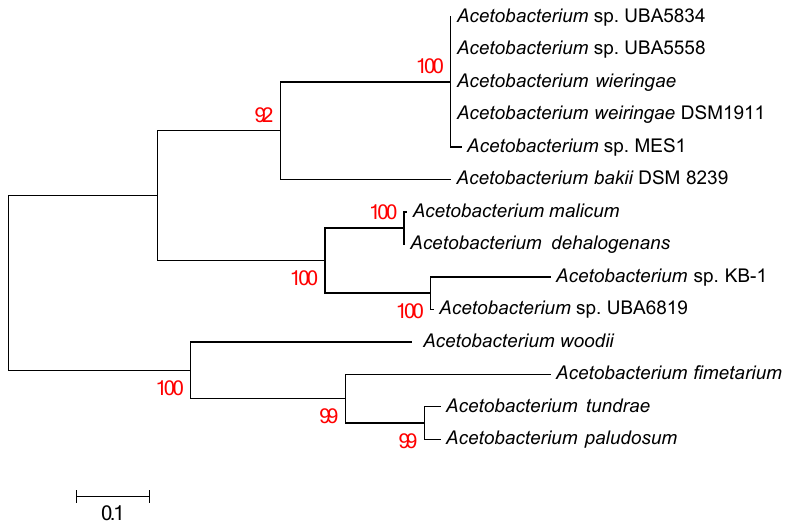

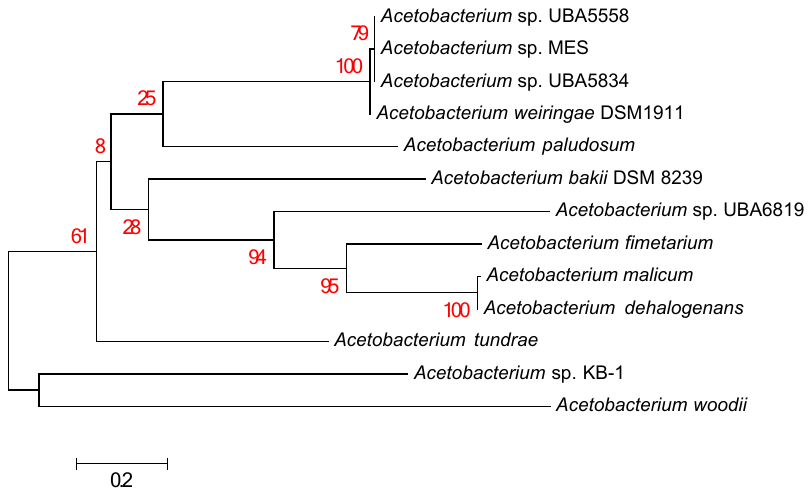

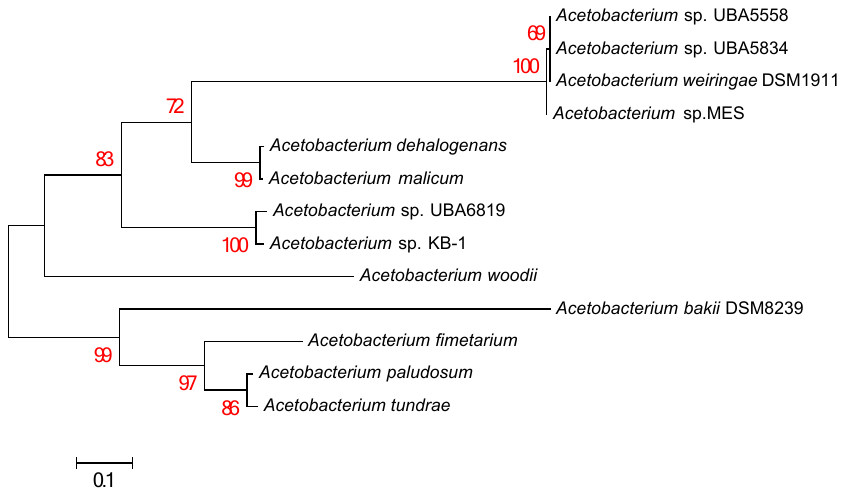

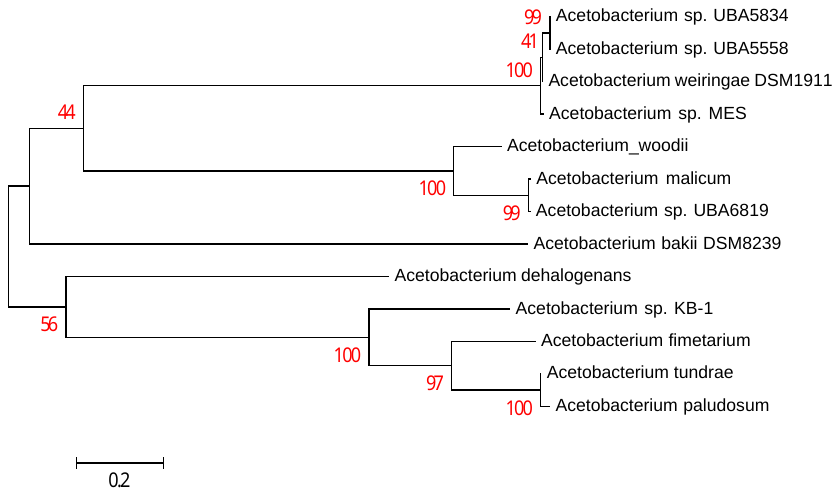

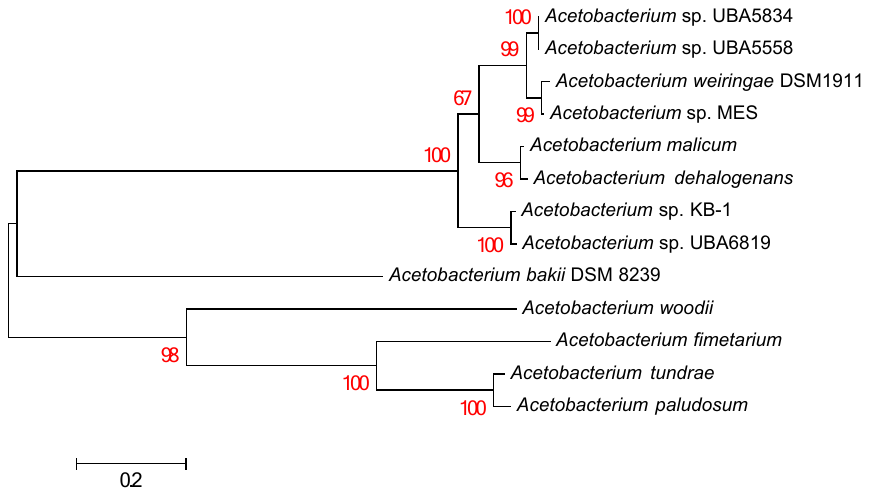

**Supplemental Figure 7. Phylogenetic analysis of 5 unique diguanylate cyclases from the *A. wieringae* clade.** Predicted protein sequences from *A.* sp. MES1 were BLASTed against each individual *Acetobacterium* genome in RAST. The top hit from each search was used to construct a maximum likelihood tree of MUSCLE-aligned protein sequences.
